# The mindful brain at rest: neural oscillations and aperiodic activity in experienced meditators

**DOI:** 10.1101/2023.10.29.564588

**Authors:** Brittany McQueen, Oscar W Murphy, Paul B Fitzgerald, Neil W. Bailey

**Author notes:** Corresponding author: Neil W Bailey.

## Abstract

**Objectives:** Previous research has demonstrated that mindfulness meditation is associated with a variety of benefits, including improved mental health. Researchers have suggested these benefits may be underpinned by differences in neural oscillations. However, previous studies measuring neural oscillations have not controlled for non-oscillatory neural activity, the power spectrum of which follows a 1/f distribution and contributes to power measurements within oscillation frequencies of interest. In this study, we applied recently developed methods to determine if past findings related to neural oscillations in meditation are present even after controlling for non-oscillatory 1/f activity.

**Methods:** 48 experienced meditators and 44 non-meditators provided resting electroencephalography (EEG) recordings. Whole scalp EEG comparisons (topographical ANOVAs) were used to test for differences between meditators and non-meditators in the distribution or global power of activity for theta, alpha, beta, and gamma oscillations, and for the 1/f components slope and intercept.

**Results:** Results indicated that meditators showed differences in theta, alpha, and gamma oscillatory power compared to non-meditators (all *p* < 0.05). Post-hoc testing suggested that the oscillatory differences were primarily driven by differences in the distribution of neural activity between meditators and non-meditators, rather than differences in the overall power across all scalp electrodes.

**Conclusion:** Our results suggest that experience with meditation is associated with higher oscillatory power and altered distributions of theta, alpha and gamma oscillations, even after controlling for non-oscillatory 1/f activity. Band-specific differences in oscillatory activity may be a mechanism through which meditation leads to neurophysiological benefits.

## Introduction

Mindfulness meditation (MM) has gained significant traction in the general population as a tool that may aid in the alleviation of daily stress and negative emotions. MM requires individuals to pay attention to the present moment with a non-judgemental awareness of the inner or outer experiences (Kabat-Zinn, 1994). Accumulative evidence suggests that MM can lead to a variety of benefits, through physiological, emotional, and cognitive changes (see Chiesa, Calati, & Serretti, 2011; Chiesa & Serretti, 2010). These benefits are supported by extensive evidence from research and meta-analyses of clinical-based interventions that target stress, depression, and anxiety (such as mindfulness-based stress reduction (MBSR) and mindfulness-based cognitive therapy (MBCT) (Chiesa & Serretti, 2010). More specifically, evidence suggests MM is associated with improvements in cognitive and attentional functioning, (Chiesa et al., 2011; Chiesa & Serretti, 2010; Delgado-Pastor, Perakakis, Subramanya, Telles, & Vila, 2013), improvements to executive functioning (Im et al., 2021), and higher-order functioning such as verbal reasoning and judgement/decision making (Gill, Renault, Campbell, Rainville, & Khoury, 2020). The benefits of MM can also be observed across facets of emotional and autonomic responses including emotional regulation and reactivity (Chiesa & Serretti, 2010; Holzel et al., 2011), improvements to autonomic regulation whilst actively engaged in meditation (Delgado-Pastor et al., 2013), and an overall reduction in automatic unconscious cognitive patterns (Burgess, Beach, & Saha, 2017; Lueke & Gibson, 2015).

Previous research has indicated that changes in the power of neural oscillations, as measured by electroencephalography (EEG), may be a neurophysiological mechanism that underpins the benefits of MM, with significant evidence in support of meditation-related differences in theta and alpha oscillatory activity (Dietl, Dirlich, Vogl, Lechner, & Strian, 1999; Kerr et al., 2011; Lomas, Ivtzan, & Fu, 2015; Sauseng, Griesmayr, Freunberger, & Klimesch, 2010). Theta activity (voltage amplitudes that cycle from negative to positive approximately 4 – 8 times per second, or at 4 – 8 Hz) has been shown to be associated with working memory functions (Klimesch, Schack, & Sauseng, 2005; Sauseng et al., 2010), anxiety, cognitive control, and decision making processes (Cavanagh & Frank, 2014; Cavanagh & Shackman, 2015) as well as attention and the processing of information (Dietl et al., 1999; Grunwald et al., 1999; Klimesch, Doppelmayr, Schimke, & Ripper, 1997). Alpha activity (voltage amplitudes that cycle from negative to positive approximately 8 – 13 times per second, or at 8 - 13 Hz) is primarily generated by parietal and occipital regions, and evidence suggests alpha activity reflects the inhibition of brain regions that are not involved in the completion of a task (Cooper, Croft, Dominey, Burgess, & Gruzelier, 2003; Jensen & Mazaheri, 2010; Klimesch, Sauseng, & Hanslmayr, 2007; Mathewson et al., 2011). Previous evidence suggests that meditators may be able to modulate alpha activity to a greater degree than non-meditators, both when selecting stimuli for attentional focus between two different sensory modalities, and when required to focus attention away from certain stimuli (Kerr et al., 2011; Wang et al., 2020). In addition, higher power for both theta and alpha oscillations in the frontal and temporal regions has been associated with a variety of types of meditation, both during meditation practice and while at rest (Kerr et al., 2011; Lee, Kulubya, Goldin, Goodarzi, & Girgis, 2018; Lomas et al., 2015; Wong, Camfield, Woods, Sarris, & Pipingas, 2015).

Less focus has been given to the study of associations between beta and gamma oscillatory activity and meditation. Beta activity (voltage amplitudes that cycle from negative to positive approximately 12 – 15 times per second, or at 12 – 25 Hz) has been linked with activation of the default mode network (DMN) – a network of neural regions involved in self-referential processing, reflective thoughts, emotional monitoring, and maladaptive rumination (Buckner, Andrews-Hanna, & Schacter, 2008; Gusnard & Raichle, 2001; Hamilton et al., 2011; Holzel et al., 2011; Jang et al., 2011; Raichle et al., 2001; Taylor et al., 2013). Specifically, a limited number of findings suggest that when compared to a resting condition, meditation is associated with decreased beta activity (Faber et al., 2015), reduced coherence of beta oscillations (as well as alpha and gamma) (Lehmann et al., 2012), and conversely greater beta amplitudes - but only for certain brain regions (Dunn, Hartigan, & Mikulas, 1999).

Gamma activity (activity above 25 Hz) has been associated with cognitive and attentional functioning, including working memory functions and the processing of sensory information (Kambara et al., 2017; Lee et al., 2018; Pritchett, Siegle, Deister, & Moore, 2015). In relation to meditation practice, greater gamma activity has been found in meditators compared to non-meditators when actively engaged in a meditative state (Lutz, Greischar, Rawlings, Ricard, & Davidson, 2004). Gamma activity has also been positively correlated with experience in meditative practices (Lee et al., 2018).

While previous research into EEG activity associated with meditation is informative, notably, many of the differences in neural activity that have been associated with meditation have been identified through investigation of neural activity during the performance of cognitive tasks, or whilst actively engaged in a meditation practice. In contrast, changes in resting-state EEG data provide novel information indicative of trait-related neural changes, specifically without the engagement of attention mechanisms that may be associated with the performance of a cognitive task or of meditation (Cahn & Polich, 2006; Lutz, Dunne, & Davidson, 2006). In contrast to the meditative state (the state of practicing meditation), meditative *traits* refer to a more persistent change, such as a shift in one’s relationship to thoughts, emotions, or a deepened sense of calmness, which may extend beyond of the period of meditation and evident in resting-state EEG analyses (Cahn & Polich, 2006; Lutz et al., 2006).

In addition, previous research using EEG to examine neural oscillations in the context of meditation has not accounted for the contribution of “non-oscillatory” neural activity (neural activity that generates voltage shifts that do not show oscillations, or a regular repeating cycle). This non-oscillatory neural activity (or “aperiodic” activity) follows a 1/f “power law”, whereby the power of neural activity at each frequency is inversely proportional to that frequency (such that as frequency increases, power decreases) (Ouyang, Hildebrandt, Schmitz, & Herrmann, 2020; Voytek et al., 2015). Recent research has shown that the 1/f activity is also important to behavioural function (Ouyang, Hildebrandt, Schmitz, & Herrmann, 2020; Voytek et al., 2015). However, the neurophysiological mechanisms that govern 1/f activity are still unclear (Ouyang et al., 2020). Most notably, traditional measures of EEG oscillatory power have not disentangled oscillatory power from 1/f non-oscillatory activity. If not addressed, this non-oscillatory 1/f activity can completely account for differences in power when measuring neural activity within specific oscillatory frequencies, such that researchers cannot draw substantiated conclusions about neural oscillations (Cahn & Polich, 2006; Kosciessa, Grandy, Garrett, & Werkle-Bergner, 2020; Lomas et al., 2015; Takahashi et al., 2005; Voytek et al., 2015). Furthermore, research measuring EEG power in lower frequency oscillations (e.g., theta (4-8 Hz) compared to gamma (>25 Hz)) is more likely to be impacted by this 1/f activity, as the amplitude of the 1/f activity in the lower frequency bands is much higher than the amplitude of the oscillatory activity (Voytek & Knight, 2015). As such, although differences in power may be detected within predetermined frequency ranges, these differences do not necessarily reflect differences in oscillatory power. The contribution of 1/f activity can thereby lead to misinterpretation of data, as there may be a number of alternative physiological processes that explain differences in frequency power measures (such as a reduction in true oscillatory power, a shift in the peak frequency of the oscillation, a reduction in power across all frequencies, or a change in the 1/f slope) (Haller et al., 2018). In consideration of this possibility, conclusions drawn by previous research regarding an association between meditation and differences in oscillatory activity may in fact be driven by non-oscillatory activity, where 1/f activity has confounded the measurement of the strength of the oscillations.

Evidence has also suggested that a flatter 1/f slope may be associated with more asynchronous neuronal firing (Voytek & Knight, 2015), as weaker synaptic inputs are associated with greater variability in synaptic input response (leading to a flatter spectrum). Alternatively, greater local positive excitatory feedback may be driven by an increased excitation/inhibition (E/I) ratio within the cortex, as higher levels of synchronous neural spiking is associated with a steeper 1/f slope (Voytek & Knight, 2015). A well-regulated E/I balance is vital for maintaining neural homeostasis, is reflective of healthy neuronal communication, and is implicated in cognitive functions such as working memory (Gao, Peterson, & Voytek, 2017; Robertson et al., 2019). Conversely, an imbalance has been associated with cognitive impairments or neurological disorders, and in certain disorders (such as schizophrenia) 1/f activity may actually be more predictive of impaired functioning than neural oscillations (Peterson, Rosen, Campbell, Belger, & Voytek, 2018). Thus, exploring potential differences in the 1/f slope and intercept may provide valuable insight into the function of 1/f activity, and comparing 1/f activity between meditators and non-meditators may indicate whether the E/I balance is modified by meditation.

Given this background, the present study utilised the extended Better Oscillation Detection (eBOSC) algorithm to identify and control for 1/f activity, in order to measure resting-state oscillatory activity from meditators and non-meditators without the confound of the 1/f activity (Kosciessa et al., 2020; Whitten, Hughes, Dickson, & Caplan, 2011). eBOSC also provides the ability to distinguish periods of EEG activity that show oscillations in specific frequencies from periods that do not show oscillations. This enables comparisons of the time spent showing oscillations above the 1/f activity in addition to oscillatory power (reflected by the measure ‘percentage of EEG trace’; see Supplementary Materials) (Haller et al., 2018; Kosciessa et al., 2020). In addition to measuring oscillatory power without 1/f activity and the slope and intercept components of the 1/f activity, the eBOSC algorithm allows analysis of peak oscillatory frequencies. Further information regarding these variables can be found in the Supplementary Materials.

The primary aims of the present study were to determine if differences in oscillatory power were present between meditators and non-meditators during resting-state EEG (whilst controlling for 1/f activity), and whether meditators and non-meditators differed in the 1/f components: slope and intercept. It was hypothesised that meditators would demonstrate greater oscillatory power for all frequency bands compared to non-meditators after controlling for 1/f activity. It was also hypothesised that there would be differences in the 1/f slope and intercept between meditators and non-meditators (reflecting differences in E/I balances). Due to the limited research examining 1/f in the context of meditation (see Rodriguez-Larios, Bracho Montes de Oca, & Alaerts, 2021), this hypothesis was non-directional. Finally, we also formed an exploratory and non-directional hypothesis that both the global amplitude of the different oscillations and 1/f parameters, and the topographical distribution of each oscillation would differ between groups, reflecting both an increase in the overall amplitude of oscillations in meditators, and a differential patterns of brain region engagement in meditators compared to non-meditators.

## Methods

### Participants

Data was collected in the context of two broader studies that examined EEG activity during cognitive tasks, which were conducted from 2016 to 2019 (results of which have already been published; see Bailey, Freedman, Raj, Spierings, et al., 2019; Bailey, Freedman, Raj, Sullivan, et al., 2019; Bailey, Raj, et al., 2019; Wang et al., 2020). Participants were recruited through community advertisement and from advertisements at meditation centres. The overall sample of resting recordings available from these two studies included 98 participants, of which 50 were experienced meditators and 48 non-meditators. Participants ranged from 19 to 64 years old. Inclusion criteria for meditators consisted of having a current meditation practice involving at least two hours per week of practice, with at least six months of meditation experience, that were mindfulness-based and met the requirements of the Kabat-Zinn definition of “paying attention in a particular way, on purpose, in the present moment, and non-judgmentally” (Kabat-Zinn, 1994). Non-meditators were only included if they had less than two hours of lifetime experience with any kind of meditation. Exclusion criteria involved self-reported current or previous mental or neurological illness, or current psychoactive medication or recreational drug use. The Beck Depression Inventory (BDI-II) and the Beck Anxiety Inventory (BAI) were administered to screen for depression and anxiety (Steer, & Beck, 1997; Beck, Steer, & Brown, 1996). For the BDI-II, participants who scored at or above the threshold for the mild range were excluded (≥19), whilst participants who scored at or above the threshold for the moderate range in the BAI were excluded (≥21). Trait mindfulness was also assessed using the Five Facet Mindfulness Questionnaire (FFMQ) (Baer, Smith, Hopkins, Krietemeyer, & Toney, 2006). Data from four non-meditators were excluded due to: insufficient EEG data (one participant), moderate anxiety (one participant), or a previous history of meditation (two participants). Data from two meditators were excluded due to a history of mental illness (one participant) or insufficient weekly meditation practice time (one participant). This left 48 meditators (27 females) and 44 non-meditators (24 females), and a total of 92 participants in the final sample (see Table 1 for participant demographics). The study was approved by the ethics committees of the Alfred Hospital and Monash University, and all participants gave written informed consent.

**Table 1:** Bayesian, Robust, and Parametric Tests for Demographic and Self-Report Data.

|  | **Meditators**<br>***M (SD)***<br>*median and MAD used*<br>*for robust tests<sup>a</sup>* | **Non-meditators**<br>***M (SD)***<br>*median and MAD used*<br>*for robust tests<sup>a</sup>* | **BF<sub>01</sub>** | **BF<sub>10</sub>** | **Statistics** |
| --- | --- | --- | --- | --- | --- |
| Sample size | 48 | 44 |  |  |  |
| Age | 37.15 (12.11) | 33 (12.98) | 1.52 | | $t(90) = -1.59, p = .116$ |
| Gender (F/M) | 27/21 | 24/20 | 3.89 | | $\chi^2(1, N = 92) = 0.03, p = .869$ |
| Years of education | 17.13 (2.43) | 16.62 (2.89) | 3.15 | | $t(89) = -0.91, p = .364$ |
| Preferred hand<br>(R/L/Ambidextrous) | 44/4/0 | 42/1/1 | 24.96 | | $\chi^2(2, N = 92) = 2.68, p = .262$ |
| BAI | 4.83 (5.35) | 5.34 (5.28) | 4.17 | | $t(90) = 0.46, p = .648$ |
| BDI-II | 0<sup>a</sup> (0<sup>a</sup>)<br>Mean 1.5 (SD = 2.54) | 0<sup>a</sup> (1<sup>a</sup>)<br>Mean 3.30 (SD = 4.70) | | 2.20 | $t(32.26^a) = 1.61^a, p = .116^a$ |
| FFMQ | 154.25 (15.4) | 134.80 (13.87) | | 1.104e+6 | $t(90) = -6.35, p < .001$ |

*Note.* Statistics refer to student's t-test (or robust Yuen's t test) and independent chi-squared tests. Robust Yuen's t tests are indicated by <sup>a</sup>. The median and MAD are reported for robust tests. MAD refers to Median absolute deviation. The mean and standard deviation (SD) are reported for parametric tests. Robust tests are based on a trimmed sample and so the degrees of freedom (if given) will not be the same for other measures. BF = Bayes Factors. BF<sub>01</sub> > 1 favours the model of the null hypothesis. BF<sub>10</sub> > 1 favours the model of the alternative hypothesis. BF<sub>01</sub> (BF<sub>excl</sub>) is reported for non-significant findings whilst BF<sub>10</sub> (BF<sub>incl</sub>) is reported for significant findings.
Higher-order interactions are excluded. All significance levels ( $\alpha$ ) set at .05. NB. The mean BDI-II score for meditators was 1.5 (SD = 2.54) and for non-meditators 3.30 (SD = 4.70) and is provided here for additional clarification as a median of 0 may not be of value to interpret on its own.

### Procedure

A Neuroscan 64-channel Ag/AgCl Quick-Cap was used to acquire EEG through NeuroScan Acquire software and a SynAmps 2 amplifier (Compumedics, Melbourne, Australia). Electrodes were referenced to an electrode between Cz and CPz, and data were collected from all electrodes for both eyes closed (EC) and eyes open (EO) conditions. Electrode impedances were kept below 5kΩ. The EEG was recorded at 1000Hz, with an online bandpass filter of 0.05 to 200Hz.

The EEG session typically lasted between 2.5 to 3.5 hours. Participants who had resting data collected during participation in the first study completed five cognitive tasks across the session, whilst participants involved in the second study completed three cognitive tasks during participation. Participants recruited from the first study completed a Go/No-Go task and a colour Stroop task prior to their resting EEG recording. Participants recruited from the second study only completed a Go/No-Go task prior to their resting EEG recording. All participants completed the eyes closed resting recording first, and then the eyes open component. Analysis of the task related data can be found in Bailey, Baell, et al. (2023), Bailey, Freedman, et al. (2019), Bailey, Geddes, et al. (2023), Bailey, Raj, et al. (2019), Payne, et al. (2019), and Wang et al. (2020). During the resting recording, participants were explicitly instructed “to rest, not to meditate”, to exert no deliberate control over their mental state, and to let their mind “do whatever it wants to”.

Pre-processing and analysis of EEG data from eyes closed and eyes open conditions were conducted for each recording separately, using the toolboxes EEGLAB (Delorme & Makeig, 2004) and FieldTrip (Oostenveld, Fries, & Maris, 2011) in MATLAB R2018b and R2020a (The MathWorks, USA). EEG data were cleaned using the RELAX cleaning pipeline (Bailey, Biabani, et al., 2022; Bailey, Hill, et al., 2022). Firstly, EEG data were filtered using a fourth order Butterworth filter with a bandpass of 1 to 80 Hz and a second order 47 to 53 Hz notch filter. The “Pre-processing” (PREP) pipeline was used to remove noisy electrodes (Bigdely-Shamlo, Mullen, Kothe, Su, & Robbins, 2015). Further remaining extreme outlying electrodes and data periods were rejected using multiple approaches as per the default settings in RELAX (full details are reported in Bailey, Biabani, et al. (2022) and Bailey, Hill, et al. (2022)). Objective artifact detection procedures were then used to identify eye movements and blinks, voltage drifts, and muscle activity. Three sequential Multiple Weiner Filter (MWF)’s were then used to filter out these artifacts and leave only clean data (Somers, Francart, & Bertrand, 2018). Following the MWF cleaning, wavelet enhanced independent component analysis was applied to reduce the artifactual components identified by ICLabel (Pion-Tonachini, Kreutz-Delgado, & Makeig, 2019) in order to clean any remaining artifacts. Further details regarding the RELAX data cleaning steps can be found in Bailey, Biabani, et al. (2022) and Bailey, Hill, et al. (2022). Lastly, the EEG data was split into epochs of 5 seconds in length with a 1.5 second overlap on each side. This provided 2 unique seconds within each epoch for frequency-power computation, and a sufficient buffer on each side of the epoch to prevent edge effects from influencing frequency-power computations. Epochs that showed improbable voltage distributions or kurtosis values exceeding 5 standard deviations for any single electrode, or 3 standard deviations for all electrodes, were rejected. In line with previous research, no more than 15% of epochs or 20% of electrodes were rejected from any participant (Bailey, Biabani et al., 2022; Kosciessa et al., 2020; see Supplementary Materials).

The eBOSC toolbox was used for the detection of theta (4-8Hz), alpha (8-12Hz), beta (12-25Hz) and gamma (>25Hz) oscillations, and the 1/f slope and intercept for each participant (Kosciessa et al., 2020; Whitten et al., 2011). The eBOSC toolbox marked periods of the EEG as containing oscillatory rhythms if the rhythmic activity exceeded a power threshold based on the aperiodic background, and if it was distinctly different from the 1/f activity within the predetermined oscillatory frequency bands. For the present study, rhythmic activity was defined as activity showing more than 3 cycles of the frequency of interest. Three cycles were specifically set as the threshold for this study, as transient oscillations between 1-3 cycles are likely to reflect a different physiological mechanism to stable (>3 cycles) oscillations (Kosciessa et al., 2020). Within the eyes closed resting-EEG recordings, 12 participants showed no periods without theta oscillations and 11 participants showed no EEG data periods without alpha oscillations. These participants were excluded from our analysis of theta and alpha power after the removal of 1/f activity, as we were unable to determine the 1/f power for these participants (and so it was not possible to subtract the 1/f activity from the oscillatory power). The mean peak frequency and percentages of the EEG trace showing oscillations for each frequency of interest were also computed by eBOSC. Further details of the eBOSC calculations and these analyses can be found in the Supplementary Materials.

### Data analysis

For statistical comparisons of each primary measure, a whole scalp analysis was conducted using the Randomisation Graphical User Interface (RAGU) (Koenig et al. 2011). RAGU allows for a combined test of potential differences in overall neural strength and distribution of activity using the Topographic Analysis of Variance (TANOVA). Separate TANOVAs were used to conduct repeated measures ANOVA design statistics examining 2 group (meditators vs non-meditators) × 2 condition (eyes closed vs eyes open) for theta, alpha, beta, and gamma oscillations after the removal of 1/f activity. These same tests were performed for the 1/f components slope and intercept. Further details regarding the application of RAGU can be found in the Supplementary Materials.

To determine whether any significant differences might be due to differences either in global power or differences in the scalp distribution of activity independent of potential differences in global power, post-hoc exploratory analyses were conducted using a Root Mean Square (RMS) test (which tests for differences in global power) and an L2 normalised TANOVA (which tests for differences in the distribution of neural activity controlling for the influence of differences in global power, by normalising RMS values within each participant to 1). Partial eta squared (ηp²) effect sizes were computed in RAGU for all comparisons of interest. Post hoc t-maps were also produced in RAGU for all measures of interest that demonstrated significant differences for each relevant TANOVA. These t-maps are equivalent to subtracting the control group data from the meditation group at every electrode. The t-maps indicate the t-values for differences at all electrodes, with the t-min value indicating where meditators showed smaller values than non-meditators, and the t-max value indicating where meditators showed larger (more positive) values than non-meditators.

Bayes Factors (BF) were also calculated in JASP for all analyses (JASP Team, 2020). BF reflect the likelihood a model can be attributed to the alternative hypothesis over the null hypothesis or vice versa (Rouder, Morey, Verhagen, Swagman, & Wagenmakers, 2017). Interaction effects were also tested for (noted by BF_incl_ / BF_excl_ values), which are calculated by comparing models that contain the effect to equivalent models stripped of the effect. BF₀₁ (and BF_excl_) is reported for non-significant findings whilst BF₁₀ (and BF_incl_) is reported for significant findings. *A priori* power analyses were not conducted, as the sample was a convenience sample obtained by combining two planned studies, and Bayesian statistics can give an indication of the confidence in null results even in the absence of a power analysis (Rouder et al., 2017).

Traditional frequentist statistical analyses were also conducted in R using the WRS2 package to implement robust statistics for mean peak frequency, RMS for all oscillations measured, and the 1/f components (Mair & Wilcox, 2020; R Core Team, 2020). For any variables which demonstrated violations in homogeneity of variance, the robust Yuen’s t-test was used instead of the independent samples t-test. As these analyses were exploratory, their full details are provided in the Supplementary Materials.

In order to control for the likelihood of increased false positive results due to the number of hypothesis tests, the Benjamini and Hochberg test (“BH”) was implemented across all primary comparisons (Benjamini & Hochberg, 1995). Primary statistical comparisons involved the main effect of group for the following variables: theta power without 1/f, alpha power without 1/f, beta power without 1/f, gamma power without 1/f, 1/f slope and 1/f intercept. Adjusted p-values are specified in the results by BH-p, and uncorrected p-values are also reported to enable comparison with previous and future research.

## Results

### Demographics and self-reported data

No differences were present between groups for any demographic variable (all p > 0.10 and BF_01_ > 1.5), with the exception of FFMQ scores (t(90) = −6.35, *p* < .001), and BDI-II scores. For BDI-II scores, robust testing (by the Yuen’s t test) did not demonstrate a significant difference (*t*(32.26) = 1.61, p = .116, BF₁₀ = 2.20) (see Table 1).

### Tests for overall differences in oscillatory power

Results from the TANOVAs without L2 normalisation demonstrated significant differences between meditators and non-meditators for the distribution or amplitude of activity in the theta, alpha, and gamma power bands (see Fig. 1-3 and Table 2 for all results, all p < .01, all BH-p < .015).

**Fig. 1.**
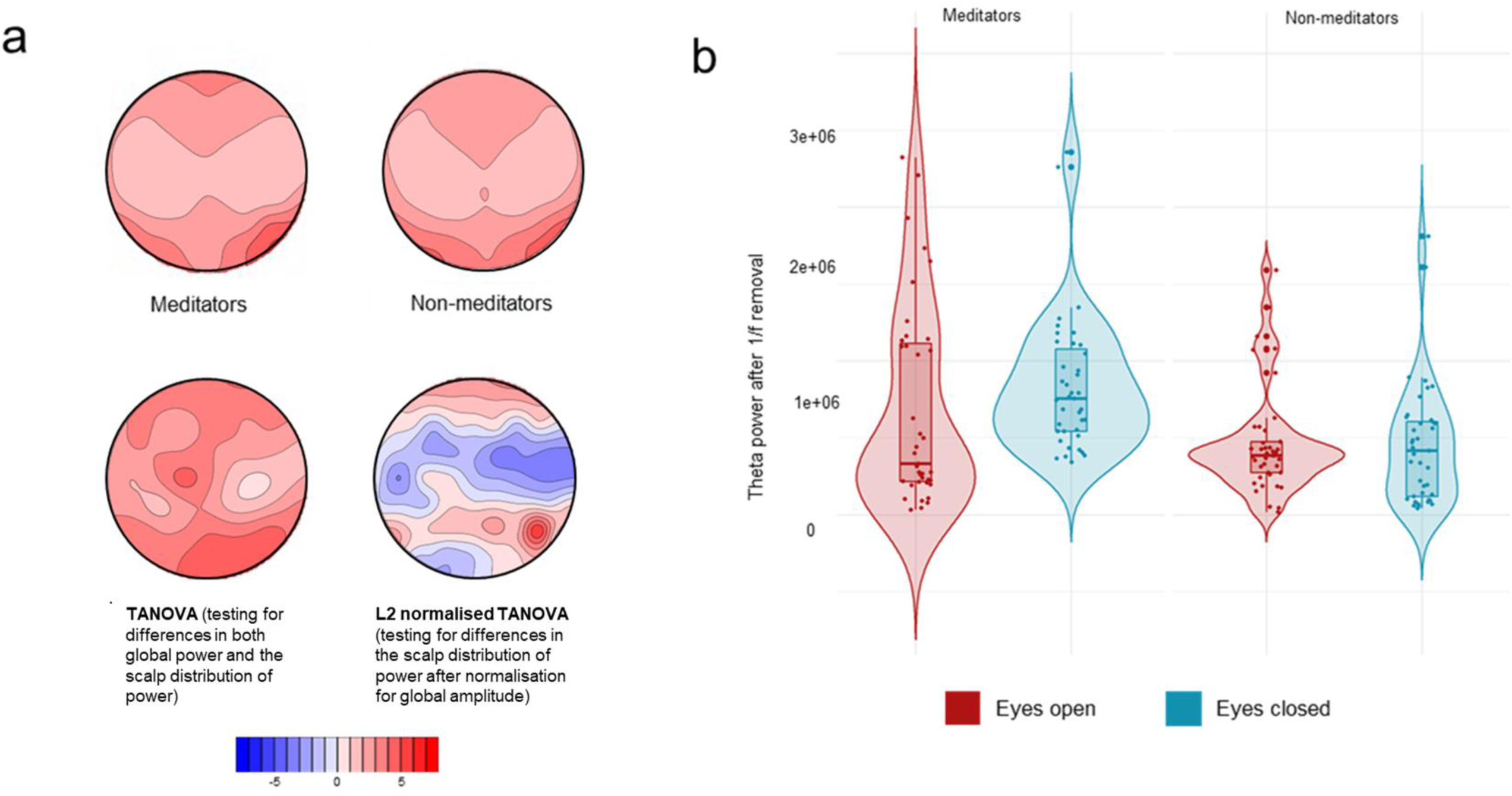
Tests for overall effect for theta power. **a** TANOVA graphs demonstrating the distribution and amplitude effect of theta power for meditators, non-meditators, the main effect (TANOVA without L2 normalisation), and the post hoc t-test (TANOVA with L2 normalisation). The blue reflects subtraction of the non-meditator group data from the meditation group data (in red). **b** Violin plots demonstrating the distribution of values for the RMS analyses of theta power for both meditators (left) and non-meditators (right), with the eyes open condition depicted in red, and eyes closed condition in blue.

**Table 2.** Statistical Results for Tests of Overall Effect for Oscillatory Power and 1/f Components.

| Variable | Effect/<br>interaction | TANOVA without L2 normalisation<br>(testing for overall effect including<br>both amplitude and distribution<br>effects) | TANOVA with L2 normalisation<br>(testing for distribution effects<br>independent of overall amplitude<br>differences) | RMS statistics |
| --- | --- | --- | --- | --- |
| Theta power (a.u.) | Main effect<br>Condition<br>Interaction | $p < .001$, BH-$p = .003^{**}$, $\eta p^2 = .139$<br>$p = .110$, BH-$p = .198$, $\eta p^2 = .029$<br>$p = .071$, BH-$p = .142$, $\eta p^2 = .036$ | $p < .001$, BFincl = $7.164e+13$, $\eta p^2 = .038$<br>$p < .001$, $\eta p^2 = .045$<br>$p < .001$, $\eta p^2 = .05$, | $p < .001$, BFincl = $52.911$, $\eta p^2 = .145$<br>$p = .313$, BFexcl = $3.650$, $\eta p^2 = .014$<br>$p = .160$, BFexcl = $1.794$, $\eta p^2 = .025$ |
| Alpha power (a.u.) | Main effect<br>Condition<br>Interaction | $p = .006$, BH-$p = .014^*$, $\eta p^2 = .066$<br>$p < .001$, BH-$p = .003^{**}$, $\eta p^2 = .532$<br>$p = .122$, BH-$p = .2$, $\eta p^2 = .057$ | $p < .001$, BFincl = $2.661e+18$, $\eta p^2 = .040$<br>$p < .001$, $\eta p^2 = .240$<br>$p = .131$, $\eta p^2 = .027$ | $p = .029$, BFincl = $1.462$, $\eta p^2 = .057$<br>$p < .001$, BFincl = $8.183e + 16$, $\eta p^2 = .591$<br>$p = .203$, BFexcl = $0.679$, $\eta p^2 = .048$ |
| Beta power (a.u.) | Main effect<br>Condition<br>Interaction | $p = .169$, BH-$p = .254$, $\eta p^2 = .030$<br>$p < .001$, BH-$p = .003^{**}$, $\eta p^2 = .499$<br>$p = .389$, BH-$p = .467$, $\eta p^2 = .022$ | $p = .061$, $\eta p^2 = .026$<br>$p < .001$, $\eta p^2 = .203$<br>$p = .062$, $\eta p^2 = .032$ | $p = .288$, BFexcl = $1.991$, $\eta p^2 = .016$<br>$p < .001$, BFincl = $3.148e + 14$, $\eta p^2 = .636$<br>$p = .947$, BFexcl = $4.212$, $\eta p^2 < .001$ |
| Gamma power (a.u.) | Main effect<br>Condition<br>Interaction | $p < .001$, BH-$p = .003^{**}$, $\eta p^2 = .191$<br>$p = .004$, BH-$p = .01^*$, $\eta p^2 = .052$<br>$p = .231$, BH-$p = .32$, $\eta p^2 = .015$ | $p < .001$, BFincl = $2.034e+25$, $\eta p^2 = .087$<br>$p = .002$, $\eta p^2 = .042$<br>$p = .033$, $\eta p^2 = .028$ | $p < .001$, BFincl = $77313.447$, $\eta p^2 = .258$<br>$p < .001$, BFincl = $4715.458$, $\eta p^2 = .234$<br>$p = .136$, BFexcl = $1.033$, $\eta p^2 = .031$ |
| Slope | Main effect<br>Condition<br>Interaction | $p = .61$, BH-$p = .66$, $\eta p^2 = .005$<br>$p < .001$, BH-$p = .003^{**}$, $\eta p^2 = .393$<br>$p = .715$, BH-$p = .715$, $\eta p^2 = .008$ | $p = .09$, $\eta p^2 = .018$<br>$p = .294$, $\eta p^2 = .013$<br>$p = .270$, $\eta p^2 = .013$ | $p = .684$, BFexcl = $2.453$, $\eta p^2 = .002$<br>$p < .001$, BFincl = $6.326e + 15$, $\eta p^2 = .585$<br>$p = .929$, BFexcl = $4.226$, $\eta p^2 < .001$ |
| Intercept | Main effect<br>Condition<br>Interaction | $p = .378$, BH-$p = .467$, $\eta p^2 = .009$<br>$p < .001$, BH-$p = .003^{**}$, $\eta p^2 = .643$<br>$p = .623$, BH-$p = .66$, $\eta p^2 = .010$ | $p = .046$, $\eta p^2 = .020$<br>$p = .025$, $\eta p^2 = .022$<br>$p = .016$, $\eta p^2 = .023$ | $p = .415$, BFexcl = $1.816$, $\eta p^2 = .007$<br>$p < .001$, BFincl = $3.142e + 24$, $\eta p^2 = .736$<br>$p = .804$, BFexcl = $4.292$, $\eta p^2 = .003$ |

Notes. Main effect compares meditators and non-meditators. Condition compares eyes closed vs eyes open conditions. Interaction compares the group (meditators and non-meditators) and condition (eyes closed and eyes open). $\eta p^2$ = Partial eta squared. a.u. = arbitrary units resulting from Morlet wavelet transform measures of power after the subtraction of the eBOSC modelled 1/f non-oscillatory activity. Adjusted p-values are specified by BH-p, and uncorrected p-values are also reported. BF = Bayes Factors. $BF_{01} > 1$ favours the model of the null hypothesis. $BF_{01}$ (BFexcl) is reported for non-significant findings whilst $BF_{10}$ (BFincl) is reported for significant findings. For the TANOVA with L2 normalisation, the BFincl value reflects a test of the interaction between Group and Electrode. RMS refers to root-mean-square. All significance levels ( $\alpha$ ) set at .05. \* $p < .05$ . \*\* $p < .01$ . \*\*\* $p < .001$ .

There was a significant difference in the pattern of neural activity between groups for theta power (p < .001, BH-p = .003, ηp² = .139). The t-map of meditator activity minus control activity for theta power was characterised by higher power in the meditator group across all electrodes, with a maximal difference at right posterior regions (t-max = 4.575 at PO6, t-min = 0.648 at C4) (Fig. 1a and 1b).

For alpha power, there was also a significant difference found between groups (p = .006, BH-p = .014, ηp² = .066) which was characterised by slightly larger values for controls over right central regions (t-min = −0.435 at C4), but larger values for meditators over other regions, with a maximal difference over central posterior regions (t-max = 5.367 at PZ) (Fig. 2a and 2b).

**Fig. 2.**
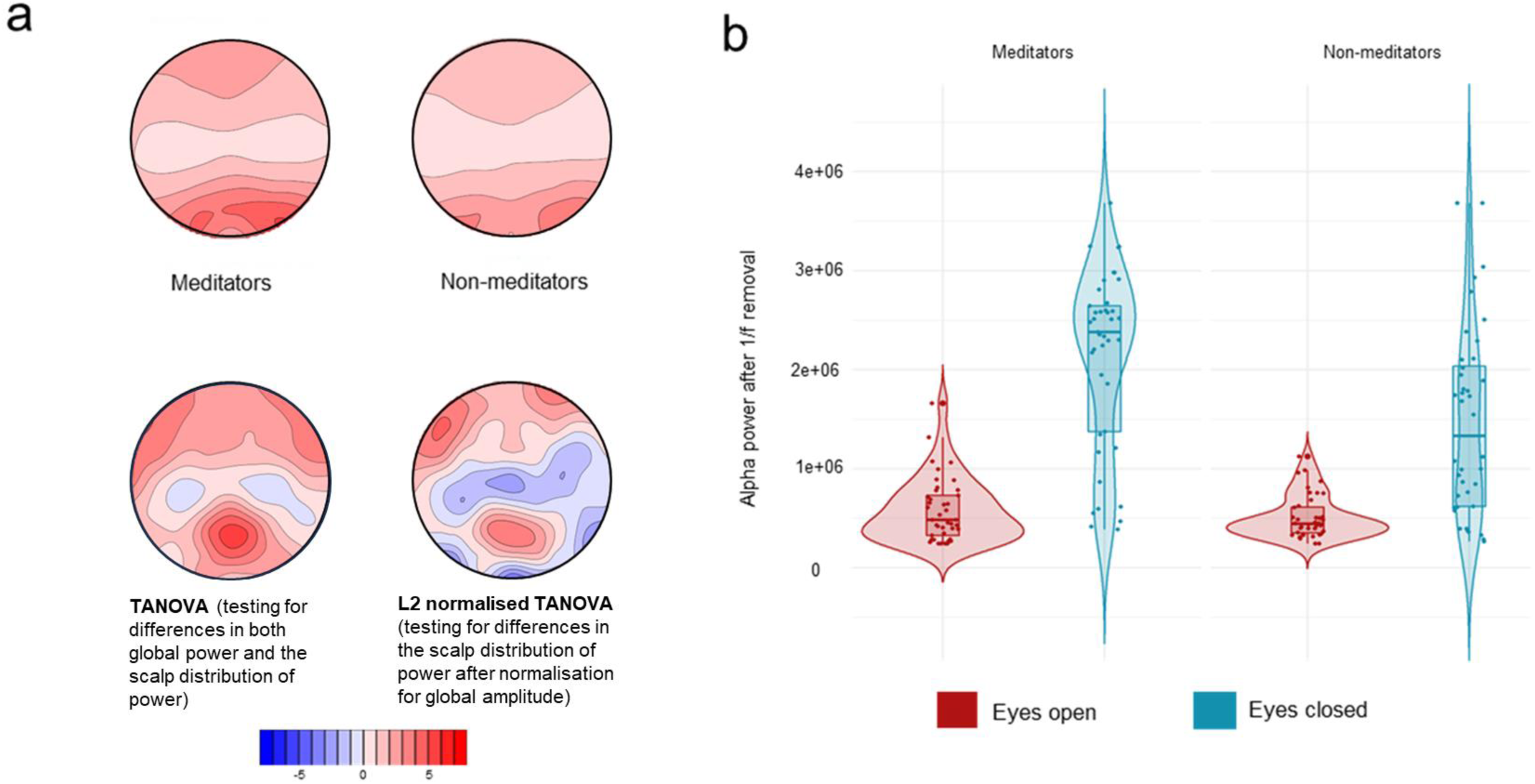
Tests for overall effect for alpha power. **a** TANOVA graphs demonstrating the distribution and amplitude effect of alpha power for meditators, non-meditators, the main effect (TANOVA without L2 normalisation), and the post hoc t-test (TANOVA with L2 normalisation). The blue reflects subtraction of the non-meditator group data from the meditation group data (in red). **b** Violin plots demonstrating the distribution of values for the RMS analyses of alpha power for both meditators (left) and non-meditators (right), with the eyes open condition depicted in red, and eyes closed condition in blue.

A significant main effect of group was also found between groups for gamma power (p < .001, BH-p = .003, ηp² = .191), and the t-map of meditator activity minus control activity was characterised by higher power values in meditators at all electrodes, with a maximal difference in middle anterior regions (t-max = 6.344 at F2, t-min = 2.912 at TP7) (Fig. 3a and 3b). However, no differences were found between groups for beta power, or the 1/f components slope and intercept (all p > .05, see Table 2), and no interactions between group and condition survived multiple comparison control (all BH-p > 0.05).

**Fig. 3.**
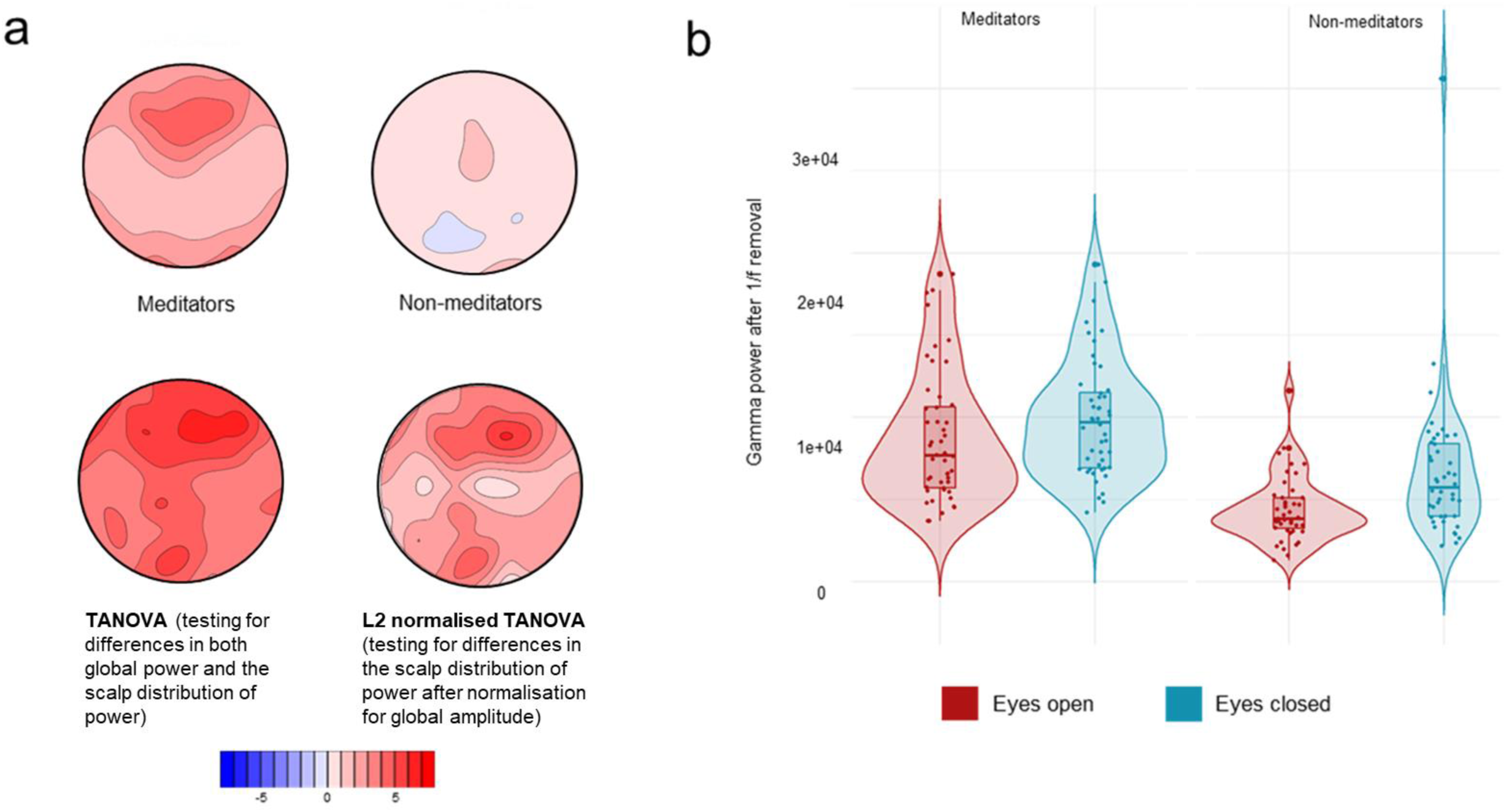
Tests for overall effect for gamma power. **a** TANOVA graphs demonstrating the distribution and amplitude effect of gamma power for meditators, non-meditators, the main effect (TANOVA without L2 normalisation), and the post hoc t-test (TANOVA with L2 normalisation). The blue reflects subtraction of the non-meditator group data from the meditation group data (in red). Note that the blue patches in the non-meditator group indicate regions that show no gamma power above the 1/f activity. **b** Violin plots demonstrating the distribution of values for the RMS analyses of gamma power for both meditators (left) and non-meditators (right), with the eyes open condition depicted in red, and eyes closed condition in blue.

As expected, significant differences were also found between conditions (eyes closed vs eyes open) for most oscillatory frequencies and 1/f components, with larger values in the eyes closed condition for the majority of variables (see Table 2, Fig. 1-3, and Supplementary Materials). Previous evidence has demonstrated that differences in oscillatory power exist between eyes closed and open conditions whilst at rest, and as such was not of primary interest to the present study and will not be explored in detail (Kirschfeld, 2005; Kosciessa et al., 2020).

### Exploratory analyses

Further exploratory and post-hoc tests were conducted to determine whether the significant results reported in our primary analyses reflected differences in the global power of the oscillation (RMS comparison) or differences in the distribution of activity across the head (reflecting a differential pattern of brain regions engaged – tested by the TANOVA with L2 normalisation).

To determine the level of support for the differences between groups in neural distribution (as demonstrated by the TANOVA with L2 normalisation), a follow up Factorial Bayesian ANOVA was conducted in JASP that included all electrodes. Prior to testing the interaction between group and electrode, the data was L2 normed and so the BF value for the interaction between group and electrodes reflects the equivalent of the normalised TANOVA (and therefore does not confound both global amplitude and distribution differences).

For theta power, both the RMS and TANOVA with L2 normalisation analyses were significant (see Table 2, 3 and 4; all p < .001), indicating significant between group differences in global power, as well as between group differences in the distribution of activity across the scalp. The t-map of meditator activity minus control activity for theta power after L2 normalisation was characterised by positive values over right posterior regions (t-max 5.971 at P6, where the meditator group showed a higher theta power) and negative values over fronto-central regions (t-min −3.541 at FC2, where the meditator group showed lower theta power values) (Fig. 1), reflecting the same pattern as the TANOVA without L2 normalisation. The effect size of the between group differences for theta power RMS was large (p < .001, ηp^²^ = .145), indicating a large between-group difference in global power. The effect size for the RMS test was larger than the effect size of the TANOVA with L2 normalisation (p < .001, ηp^²^ = .038). These combined results demonstrate the primary effect is that meditators show a higher global theta power, but also a shifted distribution of activity with meditators showing more theta in posterior regions (with a smaller effect size). These results are further depicted in Fig. 1a, and in Table 2.

**Table 3.** Statistical results for the RMS analyses of theta, alpha, and gamma power.

| **RMS analyses of theta, alpha, and gamma power** |  |  |  |  |
| --- | --- | --- | --- | --- |
|  | **Condition** | **Meditators Mdn (MAD)** | **Non-meditators Mdn (MAD)** | **Statistics** |
| Theta power (a.u) | Eyes closed | 755755.3 (357971.1) | 416944.7 (345125.5) | $p < .001$, $BF_{incl} = 52.911$, $\eta p^2 = .145$ |
|  | Eyes open | 333974.4 (344038.9) | 387094.6 (167288.2) |  |
| Alpha power (a.u) | Eyes closed | 2380160.7 (641271.7) | 1332889.2 (1062783.4) | $p = .029$, $BF_{incl} = 1.462$, $\eta p^2 = .057$ |
|  | Eyes open | 483201.4 (298702.5) | 445314.1 (157038.5) |  |
| Gamma power (a.u) | Eyes closed | 9701.58 (3664.48) | 5718.17 (2916.30) | $p < .001$, $BF_{incl} = 77313.447$, $\eta p^2 = .258$ |
|  | Eyes open | 7667.52 (3123.46) | 3817.31 (1294.39) |  |

*Notes.* RMS analyses are a measure of amplitude. $\eta p^2$ = Partial eta squared. BF = Bayes Factors. $BF_{01} > 1$ favours the model of the null hypothesis. $BF_{01}$ ( $BF_{excl}$ ) is reported for non-significant findings whilst $BF_{10}$ ( $BF_{incl}$ ) is reported for significant findings. Median and MAD were obtained from analyses conducted in R. All significance levels ( $\alpha$ ) set at .05. \* $p < .05$ . \*\* $p < .01$ . \*\*\* $p < .001$ . a.u = arbitrary units resulting from Morlet wavelet transform measures of power after the subtraction of the eBOSC modelled 1/f non-oscillatory activity. RMS refers to root-mean-square. MAD refers to Median absolute deviation.

**Table 4.**
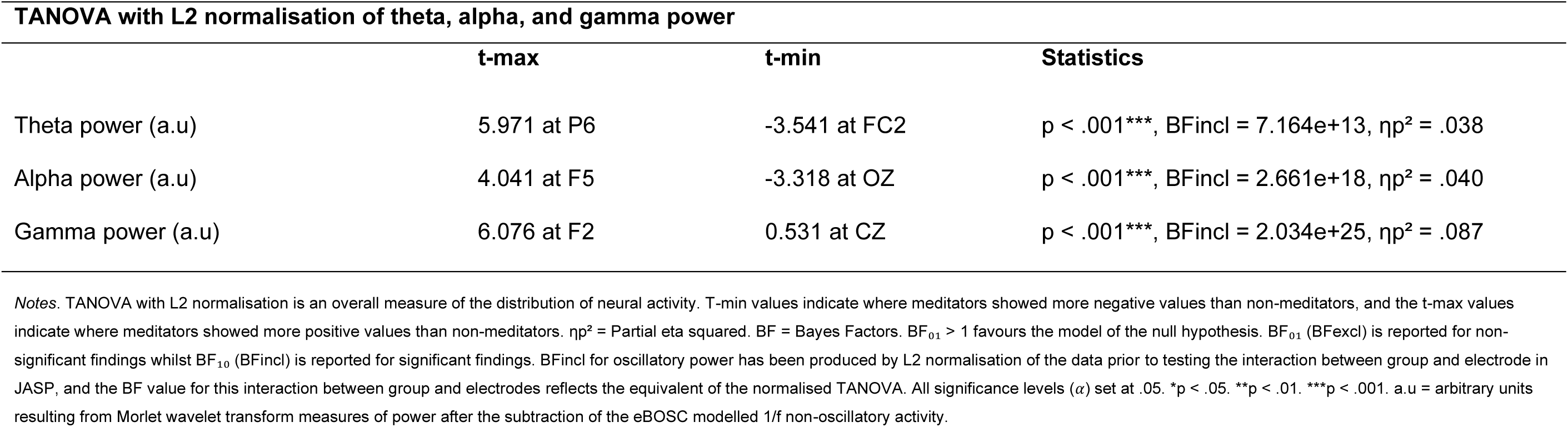
Statistical results for the TANOVA with L2 normalisation of theta, alpha and gamma power.

| **TANOVA with L2 normalisation of theta, alpha, and gamma power** |  |  |  |
| --- | --- | --- | --- |
|  | **t-max** | **t-min** | **Statistics** |
| Theta power (a.u) | 5.971 at P6 | -3.541 at FC2 | $p < .001^{***}$, $BF_{incl} = 7.164e+13$, $\eta p^2 = .038$ |
| Alpha power (a.u) | 4.041 at F5 | -3.318 at OZ | $p < .001^{***}$, $BF_{incl} = 2.661e+18$, $\eta p^2 = .040$ |
| Gamma power (a.u) | 6.076 at F2 | 0.531 at CZ | $p < .001^{***}$, $BF_{incl} = 2.034e+25$, $\eta p^2 = .087$ |

*Notes.* TANOVA with L2 normalisation is an overall measure of the distribution of neural activity. T-min values indicate where meditators showed more negative values than non-meditators, and the t-max values indicate where meditators showed more positive values than non-meditators. $\eta p^2$ = Partial eta squared. BF = Bayes Factors. $BF_{01} > 1$ favours the model of the null hypothesis. $BF_{01}$ ( $BF_{excl}$ ) is reported for non-significant findings whilst $BF_{10}$ ( $BF_{incl}$ ) is reported for significant findings. $BF_{incl}$ for oscillatory power has been produced by L2 normalisation of the data prior to testing the interaction between group and electrode in JASP, and the BF value for this interaction between group and electrodes reflects the equivalent of the normalised TANOVA. All significance levels ( $\alpha$ ) set at .05. \* $p < .05$ . \*\* $p < .01$ . \*\*\* $p < .001$ . a.u = arbitrary units resulting from Morlet wavelet transform measures of power after the subtraction of the eBOSC modelled 1/f non-oscillatory activity.

Similarly, for alpha power, both the RMS and TANOVA with L2 normalisation analyses were significant (see Table 2, 3 and 4; all p < .05). The t-map of meditator activity minus control activity for alpha power after L2 normalisation was characterised by positive values in left frontal regions (t-max 4.041 at F5, where the meditator group showed larger alpha power values) and negative values over posterior regions (t-min −3.318 at OZ, where the meditator group showed smaller alpha power values). Interestingly, these results indicate a different pattern within the topographical distribution of neural activity to the non-normalised TANOVA reported in our primary analysis, which was characterised by a maximal difference over central posterior regions (t-max 5.367 at PZ), and also indicated that meditators showed higher alpha power at all electrodes. Exploration of the L2 normalised TANOVA and RMS tests provided an explanation for this finding. Firstly, the between group effect size for the RMS analyses was small to medium (p = .029, ηp^²^ = .057), showing higher global alpha power values in the meditator group, with a larger effect size than the TANOVA with L2 normalisation (p < .001, ηp^²^ = .040). This higher power of global alpha in the meditation group drove the finding of higher posterior alpha power in the meditator group within our primary non-normalised TANOVA. However, the distribution of alpha power also differed between the groups. Within the meditator group, higher levels of frontal alpha power were present relative to overall alpha power compared to the non-meditator group. As such, when the global power differences were normalised (controlled for) and compared with the L2 normalised TANOVA, such that alpha power values over each brain region were tested relative to global alpha power within the individual, relative posterior alpha power in meditators was lower than the control group’s relative posterior alpha power. As such, the results of the L2 normalised TANOVA suggest the meditator group demonstrated an altered distribution of alpha power, with more frontal alpha produced in comparison to the control group, in addition to the increase in global alpha power. These results are depicted in Fig. 2a, and in Table 2.

For gamma power, both the RMS and TANOVA with L2 normalisation analyses were significant (see Table 2, 3 and 4; all p < .001). The t-map of meditator activity minus control activity for gamma power after L2 normalisation was characterised by positive values over fronto-central regions (t-max 6.076 at F2, where the meditator group showed higher gamma power) and negative values over central regions (t-min 0.531 at CZ, where the meditator group showed lower gamma power values) (Fig. 3), reflecting the same pattern as our primary analysis of the TANOVA without L2 normalisation. A medium to large effect size was found for the RMS analyses for gamma power (p < .001, ηp^²^ = .258), which tested for global differences in gamma power. This effect size for the RMS test was larger than the effect size of the TANOVA with L2 normalisation (p < .001, ηp^²^ = .087). The combined results from the non-normalised TANOVA and the normalised TANOVA suggest that meditators demonstrate overall higher gamma power in combination with a shifted distribution of gamma activity, with meditators showing more gamma in frontal regions relative to their global gamma power. These results are depicted in Fig. 3a, and in Table 2.

In the present study, there was also a significant main effect found for the percentage of EEG trace showing theta oscillations, with meditators demonstrating a greater percentage of the EEG trace showing oscillations within the theta frequency compared to non-meditators (value = 5.11, p = .026). These results are explored further in the Supplementary Materials.

## Discussion

The present study aimed to determine if MM is associated with differences in resting-state EEG oscillatory power after controlling for 1/f activity, and whether MM is associated with differences in the slope and intercept of 1/f non-oscillatory EEG activity. Meditators demonstrated significantly higher oscillatory power than non-meditators for theta, alpha, and gamma oscillations. The differences were driven by differences in both the distribution of activity across brain regions, and differences in the global strength of these oscillations. In particular, meditators demonstrated higher theta power as well as a shifted distribution of theta activity, with meditators demonstrating higher theta power in posterior regions relative to their global theta power. For alpha power, the meditator group demonstrated an altered distribution of activity, with higher alpha power over frontal regions (relative to the global alpha power), as well as higher global alpha power. Meditators also demonstrated an overall higher gamma power, as well as a shifted distribution of gamma power, with meditators showing higher gamma power in frontal regions relative to their global gamma power. In line with these findings, Bayesian analyses demonstrate strong support for the alternative hypothesis for group differences in both the global power and distribution of theta, alpha and gamma power. No interaction effects were observed between groups and eyes-open vs eyes-closed EEG conditions, indicating that the effects of MM were present and consistent regardless of whether participants had their eyes open or closed. No differences were found between meditators and non-meditators for the 1/f components slope and intercept, or for beta oscillatory activity. The results of the present study suggest that MM is associated with differences in oscillatory neural activity within specific frequency bands, and the use of resting-state EEG data indicates that these differences reflect trait differences, rather than simply meditation state-related differences. Furthermore, given that theta, alpha, and gamma are associated with specific cognitive functions, with larger power values typically being related to enhanced performance of those cognitive functions, the higher oscillatory power or altered distribution of oscillatory neural activity (as measured by EEG) may be one mechanism through which MM leads to benefits in cognition, attention, and general wellbeing.

### Oscillatory neural activity

#### Theta power

Within the present study, meditators demonstrated greater global theta power in comparison to non-meditators. These results mostly align with previous findings that found an association between increased theta power and meditation (Aftanas & Golocheikine, 2002; Cahn & Polich, 2006; Dunn et al., 1999; Howells, Ives-Deliperi, Horn, & Stein, 2012; Lagopoulos et al., 2009; Lomas et al., 2015; Tanaka et al., 2014). In particular, the findings of the present study are consistent with findings reported by Wong et al. (2015), whereby practiced meditators demonstrated higher theta power whilst at rest when compared to non-meditators. Importantly, the present study indicates that differences in theta activity remain significant after accounting for the potentially confounding effects of 1/f activity. These findings are of particular interest, as 1/f activity is most prominent for lower frequency ranges and thus more likely to influence the measurement of theta oscillations than other frequencies (Voytek & Knight, 2015). Additionally, given the association between theta and attentional processes (see Klimesch, Schack, & Sauseng, 2005; Sauseng et al., 2010), greater theta power may be one way through which MM leads to improvements in attention and general cognitive functioning.

#### Alpha power

For alpha power, the significant differences between meditators and non-meditators were driven by differences both in the distribution of neural activity and in global alpha power. More specifically, the topographical distribution differences were characterised by higher frontal alpha power in the meditator group (relative to global alpha power), suggesting frontal regions generated relatively greater alpha activity in meditators, in comparison to the primarily posterior alpha distribution in non-meditators. In contrast to this result, previous research has demonstrated greater alpha power in mindfulness meditators that is specific to posterior regions (Lagopoulos et al., 2009), both when comparing to a concentrative meditation practice, and separately to a relaxation condition (Dunn et al., 1999). However, this previous research did not account for the contribution of 1/f activity to measurements of alpha power, and implemented single electrode analyses which are unable to differentiate between differences in global alpha power and differences in the distribution of alpha power across the scalp. With regards to the interpretation of our alpha power results, broadly, alpha activity is thought to reflect attentional related changes, the processing of irrelevant/distracting information, and the active inhibition of processing within specific brain regions (Foxe & Snyder, 2011; Rihs, Michel, & Thut, 2009; Wang et al., 2020). Although speculative, it may be that non-meditators only inhibit posterior (visual processing related) regions whilst at rest, but continue to process memories, thoughts, and engage other attentional mechanisms, generating activity in frontal regions. Alternatively, as meditation involves focused training in attending to the sensations of the present moment, meditators may engage inhibitory mechanisms within their frontal regions, as during rest there are limited changes in sensory experience to process, and as such, the processing of non-sensory experience (for example, memories, thoughts, and attentional processes) may be reduced for practiced meditators in comparison to non-meditators.

#### Gamma power

In line with previous studies investigating gamma activity (both at rest and during tasks or meditation), the present study demonstrated that meditation experience is associated with enhanced resting gamma power (Berkovich-Ohana, Glicksohn, & Goldstein, 2011; Braboszcz, Cahn, Levy, Fernandez, & Delorme, 2017; Hauswald, Ubelacker, Leske, & Weisz, 2015; Lutz et al., 2004). Gamma activity has been associated with cognitive and attentional functioning, and higher gamma power has been associated with enhanced perceptual clarity (Kambara et al., 2017; Lee et al., 2018; Pritchett et al., 2015). There is also evidence implicating gamma activity in the role of neuroplastic changes via repetition, suggesting that increases in gamma power are positively correlated with practice (Lee et al., 2018). Greater gamma power and an altered distribution of gamma activity in the meditation group may therefore reflect a neurophysiological mechanism through which MM leads to benefits associated with cognition, attention and wellbeing, potentially reflecting the product of prolonged training of attentional processes - through a MM practice.

#### Beta power

There is some evidence that beta power increases during active meditation as compared to when at rest (Dunn, Hartigan, & Mikulas, 1999; Faber et al., 2015). However, the current study reflects the first examination of beta power during resting EEG in long-term meditators as compared to a meditation naïve group. The absence of group differences in beta power is of particular importance as it indicates that meditation was not simply associated with an overall increase in oscillatory power, but rather a selective increase in oscillatory power within certain frequency ranges. Additionally, most past research examining beta activity has not compared a meditation naïve group compared with more well practiced meditators, and instead performed comparisons between resting-state EEG and EEG recorded during active meditation practice. However, these methods only allow examination of the electrophysiological changes associated with the practice of meditation (i.e., state-dependent changes in oscillatory activity). In contrast, by comparing resting-state EEG between experienced meditators and those without a history of meditation, the current study provides valuable information regarding persistent electrophysiological changes associated with long-term meditation practice (i.e., trait-dependent changes in oscillatory activity). As general functions of beta activity include alertness, attentional arousal, and anticipatory attentional processing (Kamiński, Brzezicka, Gola, & Wróbel, 2012), it is also possible that differences between groups in these functions are found only in a state-related context, rather than during resting-state EEG (as measured by the present study). Given the limited evidence exploring beta activity and mindfulness meditators at rest, further research is needed to consolidate the findings of the present study.

### Modulation of oscillations

Overall, these accumulative findings regarding oscillatory activity provide strong evidence that long-term meditators display specific alterations in the distribution and amplitude of theta, alpha and gamma oscillations. Whilst extensive research has focussed on exploring oscillatory dynamics which underlie the meditative state, the present study demonstrates that long-term MM practice is associated with persistent changes in resting-state oscillatory activity, thereby signifying a potential neurophysiological mechanism for the long-lasting trait changes in attention and cognitive processes associated with meditation practice. In addition, rather than just an overall difference in the strength of oscillations, the present study highlights that differences in the topographical distribution of activity (reflecting altered engagement of brain regions) for these oscillations drive these results, indicating that meditation may lead to differential engagement of neural activity.

The differences observed in the topographical distribution of oscillations and/or amplitude of different oscillatory patterns may be a result of the modulation of neural activity by meditators, dependent on the requirements at hand. Modulation of neural signals contributes to healthy cognition (Armbruster-Genç, Ueltzhöffer, & Fiebach, 2016) through helping us adapt in times of uncertainty (Kosciessa, Lindenberger, & Garrett, 2021), and adapt to our environment (Kloosterman, Kosciessa, Lindenberger, Fahrenfort, & Garrett, 2020). Previous research has demonstrated that experienced meditators display a greater ability for the modulation of oscillations, specifically for theta and alpha bands (Tanaka et al., 2014; Wang et al., 2020). Wang et al. (2020) found meditators demonstrated a greater ability to modulate alpha distribution between low task demands and high task demands - which may require more neural resources. In line with these past findings, the present study may not necessarily reflect that meditation consistently results in larger oscillatory power across theta, alpha, gamma and beta power. It may be that meditators are able to modulate neural activity with either increases or decreases in favour of task-relevant processing regions, leading to a larger range for modulation, and a MM-related increase in one’s ability to respond to their environment. Changes that result from meditation may therefore reflect enhancement not of one specific neural process, but of the modulation of a range of oscillatory activity which support cognitive, emotional or attentional processing. However, we note that that our recent “highly comparative” analysis that assessed a massive number of statistical properties of the EEG time-series showed that features related to the stationarity of the EEG data (the consistency of statistical properties across different periods within the EEG data) provided more successful classification of meditators than oscillatory measures (Bailey, Fulcher, Caldwell, Hill, Fitzgibbon, van Dijk, & Fitzgerald, 2023). However, band power measures assessed in that study did not account for 1/f activity, and only assessed band power within the top eight principal components of the EEG data, and within a small number of single electrodes. As such, the current study more comprehensively characterises differences in oscillatory activity between meditators and non-meditators (Bailey, Fulcher, Caldwell, Hill, Fitzgibbon, van Dijk, & Fitzgerald, 2023).

### Predictive coding and free energy principle

Our findings may provide further insight when viewed within the conceptual framework of the free energy principle (FEP). The FEP suggests that the brain maximizes efficiency through proactive and anticipatory modelling of its environment, and thereby minimises ‘free-energy’ arising from prediction errors and the likelihood of ‘surprise’ (Friston, 2013). Through construction of hierarchical predictive models that have been selected through processes analogous to Bayesian probabilistic reasoning (e.g., based on model fit and prior beliefs), the brain (and nervous system) functions by minimizing prediction error (the mismatch between its prior model and incoming sensory information). Within this free-energy minimization framework, models constructed by the brain can be updated to better fit the world (perceptual inference) or update the world (through motor control of the body’s musculature) to better fit the brain’s prior model (active inference). We propose two theoretical perspectives for how mindfulness may affect parameters of the predictive coding framework that align with our results. Firstly, the increase in theta and gamma power shown in our results might reflect “fact free learning”. Fact free learning has been suggested to occur in the brain, whereby the brain constructs better models that have greater explanatory power through iterative adjustments of existing priors, without additional sensory information (priors) or active inference (taking action to ensure sensory information aligns with the brain’s model) (Friston et al., 2017). Secondly, meditators might show a reduction in counterfactual processing, which may align with the shift towards alpha power with a more frontal distribution in meditators, where counterfactual processing taking place in the higher regions of the predictive coding hierarchy (reflected by the frontal regions) might be inhibited. “Counterfactual” processing refers to the modelling (or internal simulation) of sensory states that an individual may observe if they were to perform or participate in actions under a particular set of model parameters (e.g., possible outcomes) (Corcoran, Pezzulo, & Hohwy, 2020). This allows for the evaluation of the expected prediction error (free energy) from a variety of actions under alternative contexts before making a decision and taking action.

Laukkonen and Slagter (2021) suggest that meditation may reduce “counterfactual” temporally deep cognition and reduce predictive abstraction through being in the “here and now”, leading to greater flexibility in daily life. We note that these two explanations could be seen as conflicting, and furthermore that the finding of increased theta and gamma power concurrent with increased alpha power (which is thought to reflect an inhibition of activity within a brain region) might also be seen to be a conflicting finding. Further research is required to determine the functional relevance and the physiological explanation of these findings.

### 1/f and excitation/inhibition balance

Previous research has suggested that differences in the 1/f slope may reflect differences in the E/I balance in the brain (Haller et al., 2018; Voytek & Knight, 2015). In relation to meditation, one study has reported differences between meditators and novices in the 1/f slope, with experienced meditators demonstrating a more negative (steeper) slope during meditation relative to rest, and novice meditators presenting the opposite pattern – a flatter slope during meditation relative to rest (Rodriguez-Larios, Bracho Montes de Oca, & Alaerts, 2021). Rodriguez-Larios, Bracho Montes de Oca, & Alaerts, (2021) have suggested these findings may be due to the fact that novices found the meditation condition more cognitively demanding than rest (leading to a flatter slope and a higher E/I ratio), whilst the opposite was true for more experienced meditators (leading to a steeper 1/f slope and lower E/I ratio).

Contrary to our hypotheses and prior research, no differences were found between meditators and non-meditators for 1/f slope or intercept in the present study. This result suggests that whilst differences in oscillatory activity are present, meditation is not associated with differences in neural activity produced by altered E/I balances related to neuroplastic change (primarily modulated by GABAergic and glutamatergic neurotransmitters). These findings are particularly interesting given that 1/f activity has also been found to be functionally relevant for perceptual processing (such as perceptual decision-making), visuomotor performance (Immink et al., 2021), and may also be more reflective of cognitive performance in comparison to the measure of oscillatory activity (Peterson et al., 2018). The null findings and large BF01 values of the present study indicate that differences in the E/I balance of the brain at rest are unlikely to be related to long-term meditation practice, and may not be the explanatory mechanism for improved attention, mental health, and wellbeing from MM. It is worth noting however, that because our study focused on healthy participants, the current results cannot rule out the possibility that meditation may lead to improvements in E/I balances found in clinical populations where E/I balances may be atypical prior to a meditation intervention (Peterson et al., 2018). Rather, it may be that the E/I balance is not altered by meditation when it is already functioning adequately in healthy controls. In addition, this null finding also provides important validation for previous studies demonstrating that MM is associated with differences in oscillatory power. Given that 1/f activity can significantly influence the measurement of oscillatory power if not properly controlled, we can be confident that the observed differences in theta, alpha, and gamma power in the present study do indeed reflect changes in long-term MM rather than simply reflecting 1/f changes.

### Limitations and future directions

Although significant differences were found between meditators and non-meditators for theta, alpha and gamma activity, the findings of the present study were cross-sectional, and therefore causation cannot be determined. Additionally, while the meditator and non-meditator groups in the current study were matched on key demographic variables which may influence oscillatory activity (such as age and gender), it is also possible that individuals that practice meditation have different lifestyle habits or participate in other activities not accounted for that may influence changes in neural activity (e.g. diet, exercise, and substance use; see Cramer, Sibbritt, Park, Adams, & Lauche (2017)). Future research could control for this by exploring what other lifestyle habits participants may be involved in and performing between/within group comparisons to determine if significant differences are present. Future research would also benefit from testing causal relationships through longitudinal studies (although these studies are exceedingly difficult to achieve in practice given the “long-term” nature of meditation practice).

Additionally, our results may be specific to the conditions of the present study, which recruited healthy participants across a broad age range, and who practiced MM (with recruitment not constrained to specific sub-types of a MM practice). Further research should be conducted to explore whether these trait changes are consistent with different populations, such as different age groups or alternative meditation sub-groups.

As meditation practices are a subjective endeavour, the number of hours an individual has invested in meditation may not necessarily equate to how advanced someone is in their practice. The inclusion criteria of our study required experienced meditators to have a minimum of 6 months of consistent practice – however one meditator had 48 years of experience. Even though the number of years individuals practice for may be considered a subjective account for being an experienced meditator, certain benefits may be associated with length of time spent meditating. Our study did confirm that the groups differed significantly in trait mindfulness as measured by the FFMQ, which was expected given that this measure may be influenced by previous meditation experience (Pang & Ruch, 2019). However, to further explore this issue, future research would benefit from recruiting participants specifically from distinct categories of experience (for example, novice, somewhat experienced, and very experienced meditators), or a larger sample size that could provide sufficient statistical power to robustly assess correlations between experience and oscillatory power.

Finally, whilst participants were under explicit instruction to refrain from meditating during the resting EEG recordings, and to rest without any deliberate control over their mental contents, we cannot verify that they followed this instruction and were not meditating. Even if participants were following these instructions, it may be that a meditative state of mind is a long-term meditator’s typical baseline resting state, which presents the possibility that different mental states were measured between the two groups. However, whilst this may be a critique of the current study, we note that it is a limitation that is impossible to address, since beyond a participant’s self-report, there is no way to verify whether a participant is in the state of rest or meditation. Additionally, even if the meditator’s baseline resting state is more similar to a meditative state than a typical non-meditator’s resting state, we feel that our results are still informative about neural oscillations in long-term meditators during “resting” periods, as this limitation indirectly suggests that a meditator’s resting state throughout the day contains brain activity that is more similar to the meditative state. Moreover, one of the primary motivations for adopting a MM practice is that beneficial physical and psychological effects persist outside of the meditation practice itself, hence the need for studies examining the neural correlates of MM in a resting-state reflected by the present study.

## Ethics Information

Ethics approval was provided by the Ethics Committees of Monash University and Alfred Hospital. All participants provided written informed consent prior to participation in the study.

## Acknowledgements

We gratefully acknowledge the Dhamma Aloka Vipassana meditation centre in Melbourne and the Melbourne Insight Meditation centre for their assistance with the recruitment of several meditators who took part in the study.

## References

Aftanas, L. I., & Golocheikine, S. A. (2002). Non-linear dynamic complexity of the human EEG during meditation. Neuroscience Letters, 330(2), 143–146. doi:10.1016/S0304-3940(02)00745-0

Armbruster-Genç, D. J. N., Ueltzhöffer, K., & Fiebach, C. J. (2016). Brain Signal Variability Differentially Affects Cognitive Flexibility and Cognitive Stability. The Journal of Neuroscience, 36(14), 3978–3987. doi:10.1523/jneurosci.2517-14.2016

Baer, R. A., Smith, G. T., Hopkins, J., Krietemeyer, J., & Toney, L. (2006). Using self-report assessment methods to explore facets of mindfulness. Assessment, 13(1), 27–45.

Bailey, N. W., Baell, O., Payne, J. E., Humble, G., Geddes, H., Cahill, I., & Fitzgerald, P. B. (2023). Experienced Meditators Show Multifaceted Attention-Related Differences in Neural Activity. Mindfulness, 1–29.

Bailey, N. W., Biabani, M., Hill, A. T., Miljevic, A., Rogasch, N. C., McQueen, B., … Fitzgerald, P. B. (2023). Introducing RELAX: An automated pre-processing pipeline for cleaning EEG data-Part 1: Algorithm and application to oscillations. Clinical Neurophysiology, 149, 178–201.

Bailey, N. W., Hill, A. T., Biabani, M., Murphy, O. W., Rogasch, N. C., McQueen, B., … Fitzgerald, P. B. (2023). RELAX part 2: A fully automated EEG data cleaning algorithm that is applicable to Event-Related-Potentials. Clinical Neurophysiology, 149, 202–222.

Bailey, N. W., Freedman, G., Raj, K., Spierings, K. N., Piccoli, L., Sullivan, C., … Fitzgerald, P. (2019). Mindfulness meditators show enhanced working memory performance concurrent with different brain region engagement patterns during recall. BioRxiv, 801746. doi:10.1101/801746

Bailey, N. W., Fulcher, B. D., Caldwell, B., Hill, A. T., Fitzgibbon, B., van Dijk, H., & Fitzgerald, P. B. (2023). Uncovering a stability signature of brain dynamics associated with meditation experience using massive time-series feature extraction. bioRxiv, 2023–06.

Bailey, N. W., Geddes, H., Zannettino, I., Humble, G., Payne, J., Baell, O., … Fitzgerald, P. B. (2023). Meditators probably show increased behaviour-monitoring related neural activity. Mindfulness, 14(1), 33–49.

Bailey, N. W., Freedman, G., Raj, K., Spierings, K. N., Piccoli, L. R., Sullivan, C. M., … Fitzgerald, P. B. (2020). Mindfulness Meditators Show Enhanced Accuracy and Different Neural Activity During Working Memory. Mindfulness, 11(7), 1762–1781. doi:10.1007/s12671-020-01393-8

Bailey, N. W., Freedman, G., Raj, K., Sullivan, C. M., Rogasch, N. C., Chung, S. W., … Fitzgerald, P. B. (2019). Mindfulness meditators show altered distributions of early and late neural activity markers of attention in a response inhibition task. PLoS One, 14(8), e0203096. doi:10.1371/journal.pone.0203096

Beck, A. T., Steer, R. A., & Brown, G. K. (1996). Beck Depression Inventory-II. San Antonio, 78(2), 490–498.

Benjamini, Y., and Hochberg, Y. (1995). Controlling the false discovery rate: a practical and powerful approach to multiple testing. Journal of the Royal Statistical Society Series B, 57, 289–300. http://www.jstor.org/stable/2346101.

Berkovich-Ohana, A., Glicksohn, J., & Goldstein, A. (2011). Mindfulness-induced changes in gamma band activity – Implications for the default mode network, self-reference and attention. Clin Neurophysiol, 123(4), 700–710. doi:10.1016/j.clinph.2011.07.048

Bigdely-Shamlo, N., Mullen, T., Kothe, C., Su, K.-M., & Robbins, K. A. (2015). The PREP pipeline: standardized preprocessing for large-scale EEG analysis. Frontiers in Neuroinformatics, 9(16). doi:10.3389/fninf.2015.00016

Braboszcz, C., Cahn, B. R., Levy, J., Fernandez, M., & Delorme, A. (2017). Increased Gamma Brainwave Amplitude Compared to Control in Three Different Meditation Traditions. PLoS One, 12(1), e0170647–e0170647. doi:10.1371/journal.pone.0170647

Buckner, R. L., Andrews-Hanna, J. R., & Schacter, D. L. (2008). The brain’s default network: anatomy, function, and relevance to disease. Ann N Y Acad Sci, 1124, 1–38. doi:10.1196/annals.1440.011

Burgess, D. J., Beach, M. C., & Saha, S. (2017). Mindfulness practice: A promising approach to reducing the effects of clinician implicit bias on patients. Patient Education and Counseling, 100(2), 372–376. doi:10.1016/j.pec.2016.09.005

Cahn, B. R., & Polich, J. (2006). Meditation states and traits: EEG, ERP, and neuroimaging studies. Psychol Bull, 132(2), 180–211. doi:10.1037/0033-2909.132.2.180

Cavanagh, J. F., & Frank, M. J. (2014). Frontal theta as a mechanism for cognitive control. Trends in Cognitive Sciences, 18(8), 414–421. doi:10.1016/j.tics.2014.04.012

Cavanagh, J. F., & Shackman, A. J. (2015). Frontal midline theta reflects anxiety and cognitive control: Meta-analytic evidence. Journal of Physiology - Paris, 109(1-3), 3–15. doi:10.1016/j.jphysparis.2014.04.003

Chiesa, A., Calati, R., & Serretti, A. (2011). Does mindfulness training improve cognitive abilities? A systematic review of neuropsychological findings. Clinical Psychology Review, 31(3), 449–464. doi:10.1016/j.cpr.2010.11.003

Chiesa, A., & Serretti, A. (2010). A systematic review of neurobiological and clinical features of mindfulness meditations. Psychol Med, 40(8), 1239–1252. doi:10.1017/S0033291709991747

Cooper, N. R., Croft, R. J., Dominey, S. J. J., Burgess, A. P., & Gruzelier, J. H. (2003). Paradox lost? Exploring the role of alpha oscillations during externally vs. internally directed attention and the implications for idling and inhibition hypotheses. International Journal of Psychophysiology, 47(1), 65–74. doi:10.1016/S0167-8760(02)00107-1

Corcoran, A. W., Pezzulo, G., & Hohwy, J. (2020). From allostatic agents to counterfactual cognisers: active inference, biological regulation, and the origins of cognition. Biology & Philosophy, 35(3). doi:10.1007/s10539-020-09746-2

Cramer, H., Sibbritt, D., Park, C. L., Adams, J., & Lauche, R. (2017). Is the practice of yoga or meditation associated with a healthy lifestyle? Results of a national cross-sectional survey of 28,695 Australian women. J Psychosom Res, 101, 104–109. doi:10.1016/j.jpsychores.2017.07.013

Delgado-Pastor, L. C., Perakakis, P., Subramanya, P., Telles, S., & Vila, J. (2013). Mindfulness (Vipassana) meditation: effects on P3b event-related potential and heart rate variability. Int J Psychophysiol, 90(2), 207–214. doi:10.1016/j.ijpsycho.2013.07.006

Delorme, A., & Makeig, S. (2004). EEGLAB: an open source toolbox for analysis of single-trial EEG dynamics including independent component analysis. Journal of neuroscience methods, 134(1), 9–21. doi:10.1016/j.jneumeth.2003.10.009

Dietl, T., Dirlich, G., Vogl, L., Lechner, C., & Strian, F. (1999). Orienting response and frontal midline theta activity: a somatosensory spectral perturbation study. Clinical Neurophysiology, 110, 1204–1209.

Dunn, B., Hartigan, J., & Mikulas, W. (1999). Concentration and Mindfulness Meditations: Unique Forms of Consciousness? Applied Psychophysiology and Biofeedback, 24(3), 147–165. doi:10.1023/A:1023498629385

Faber, P. L., Lehmann, D., Gianotti, L. R. R., Milz, P., Pascual-Marqui, R. D., Held, M., & Kochi, K. (2015). Zazen meditation and no-task resting EEG compared with LORETA intracortical source localization. Cognitive Processing, 16(1), 87–96. doi:10.1007/s10339-014-0637-x

Foxe, J., & Snyder, A. (2011). The Role of Alpha-Band Brain Oscillations as a Sensory Suppression Mechanism during Selective Attention. Frontiers in psychology, 2(154). doi:10.3389/fpsyg.2011.00154

Friston, K. (2013). Life as we know it. Journal of The Royal Society Interface, 10(86), 20130475. doi:10.1098/rsif.2013.0475

Friston, K., Lin, M., Frith, C., Pezzulo, G., Hobson, J., & Ondobaka, S. (2017). Active Inference, Curiosity and Insight. Neural Computation, 29, 1–51. doi:10.1162/neco_a_00999

Gao, R., Peterson, E., & Voytek, B. (2017). Inferring synaptic excitation/inhibition balance from field potentials. Neuroimage, 158, 70. doi:10.1016/j.neuroimage.2017.06.078

Gill, L.-N., Renault, R., Campbell, E., Rainville, P., & Khoury, B. (2020). Mindfulness induction and cognition: A systematic review and meta-analysis. Consciousness and Cognition, 84, 102991. doi:10.1016/j.concog.2020.102991

Grunwald, M., Weiss, T., Krause, W., Beyer, L., Rost, R., Gutberlet, I., & Gertz, H.-J. (1999). Power of theta waves in the EEG of human subjects increases during recall of haptic information. Neuroscience Letters, 260, 189–192.

Gusnard, D. A., & Raichle, M. E. (2001). Searching for a baseline: Functional imaging and the resting human brain. Nat Rev Neurosci, 2.

Haller, M., Donoghue, T., Peterson, E., Varma, P., Gao, R., Noto, T., … Voytek, B. (2018). Parameterizing neural power spectra. BioRxiv. doi:10.1101/299859

Hamilton, J. P., Furman, D. J., Chang, C., Thomason, M. E., Dennis, E., & Gotlib, I. H. (2011). Default-Mode and Task-Positive Network Activity in Major Depressive Disorder: Implications for Adaptive and Maladaptive Rumination. Biological Psychiatry, 70(4), 327–333. doi:10.1016/j.biopsych.2011.02.003

Hauswald, A., Ubelacker, T., Leske, S., & Weisz, N. (2015). What it means to be Zen: marked modulations of local and interareal synchronization during open monitoring meditation. Neuroimage, 108, 265–273. doi:10.1016/j.neuroimage.2014.12.065

Holzel, B. K., Lazar, S. W., Gard, T., Schuman-Olivier, Z., Vago, D. R., & Ott, U. (2011). How Does Mindfulness Meditation Work? Proposing Mechanisms of Action From a Conceptual and Neural Perspective. Perspect Psychol Sci, 6(6), 537–559. doi:10.1177/1745691611419671

Howells, F. M., Ives-Deliperi, V. L., Horn, N. R., & Stein, D. J. (2012). Mindfulness based cognitive therapy improves frontal control in bipolar disorder: a pilot EEG study. BMC psychiatry, 12(1), 15. doi:10.1186/1471-244X-12-15

Im, S., Stavas, J., Lee, J., Mir, Z., Hazlett-Stevens, H., & Caplovitz, G. (2021). Does mindfulness-based intervention improve cognitive function?: A meta-analysis of controlled studies. Clinical Psychology Review, 84, 101972. doi:10.1016/j.cpr.2021.101972

Immink, M. A., Cross, Z. R., Chatburn, A., Baumeister, J., Schlesewsky, M., & Bornkessel-Schlesewsky, I. (2021). Resting-state aperiodic neural dynamics predict individual differences in visuomotor performance and learning. BioRxiv, 2021.2004.2008.438941. doi:10.1101/2021.04.08.438941

Jang, J. H., Jung, W. H., Kang, D. H., Byun, M. S., Kwon, S. J., Choi, C. H., & Kwon, J. S. (2011). Increased default mode network connectivity associated with meditation. Neurosci Lett, 487(3), 358–362. doi:10.1016/j.neulet.2010.10.056

Jensen, O., & Mazaheri, A. (2010). Shaping functional architecture by oscillatory alpha activity: gating by inhibition. Frontiers in human neuroscience, 4, 186. doi:10.3389/fnhum.2010.00186

Kabat-Zinn, J. (1994). Wherever You Go, There You Are: Mindfulness Meditation in Everyday Life. In (Vol. 119, pp. 124). New York: Media Source.

Kambara, T., Brown, E. C., Jeong, J.-W., Ofen, N., Nakai, Y., & Asano, E. (2017). Spatio-temporal dynamics of working memory maintenance and scanning of verbal information. Clinical Neurophysiology, 128(6), 882–891. doi:10.1016/j.clinph.2017.03.005

Kamiński, J., Brzezicka, A., Gola, M., & Wróbel, A. (2012). Beta band oscillations engagement in human alertness process. International Journal of Psychophysiology, 85(1), 125–128. doi:10.1016/j.ijpsycho.2011.11.006

Kerr, C. E., Jones, S. R., Wan, Q., Pritchett, D. L., Wasserman, R. H., Wexler, A., … Moore, C. I. (2011). Effects of mindfulness meditation training on anticipatory alpha modulation in primary somatosensory cortex. Brain Research Bulletin, 85(3-4), 96–103. doi:10.1016/j.brainresbull.2011.03.026

Kirschfeld, K. (2005). The physical basis of alpha waves in the electroencephalogram and the origin of the “Berger effect”. Biological Cybernetics, 92(3), 177–185. doi:10.1007/s00422-005-0547-1

Klimesch, W., Doppelmayr, M., Schimke, H., & Ripper, B. (1997). Theta synchronization and alpha desynchronization in a memory task. Psychophysiology, 34.

Klimesch, W., Sauseng, P., & Hanslmayr, S. (2007). EEG alpha oscillations: The inhibition– timing hypothesis. Brain Research Reviews, 53(1), 63–88. doi:10.1016/j.brainresrev.2006.06.003

Klimesch, W., Schack, B., & Sauseng, P. (2005). The Functional Significance of Theta and Upper Alpha Oscillations. Experimental Psychology, 52(2), 99–108. doi:10.1027/1618-3169.52.2.99

Kloosterman, N. A., Kosciessa, J. Q., Lindenberger, U., Fahrenfort, J. J., & Garrett, D. D. (2020). Boosts in brain signal variability track liberal shifts in decision bias. eLife, 9. doi:10.7554/elife.54201

Koenig, T., Kottlow, M., Stein, M., & Melie-García, L. (2011). Ragu: A Free Tool for the Analysis of EEG and MEG Event-Related Scalp Field Data Using Global Randomization Statistics. Computational Intelligence and Neuroscience, 2011, 1–14. doi:10.1155/2011/938925

Kosciessa, J. Q., Grandy, T. H., Garrett, D. D., & Werkle-Bergner, M. (2020). Single-trial characterization of neural rhythms: Potential and challenges. Neuroimage, 206, 116331. doi:10.1016/j.neuroimage.2019.116331

Kosciessa, J. Q., Lindenberger, U., & Garrett, D. D. (2021). Thalamocortical excitability modulation guides human perception under uncertainty. Nature Communications, 12(1). doi:10.1038/s41467-021-22511-7

Lagopoulos, J., Xu, J., Rasmussen, I., Vik, A., Malhi, G., Eliassen, C., … Ellingsen, Ø. (2009). Increased Theta and Alpha EEG Activity During Nondirective Meditation. Journal of alternative and complementary medicine (New York, N.Y.), 15, 1187–1192. doi:10.1089/acm.2009.0113

Laukkonen, R. E., & Slagter, H. A. (2021). From many to (n)one: Meditation and the plasticity of the predictive mind. Neuroscience & Biobehavioral Reviews, 128, 199–217. doi:10.1016/j.neubiorev.2021.06.021

Lee, D. J., Kulubya, E., Goldin, P., Goodarzi, A., & Girgis, F. (2018). Review of the Neural Oscillations Underlying Meditation. Front Neurosci, 12, 178. doi:10.3389/fnins.2018.00178

Lehmann, D., Faber, P. L., Tei, S., Pascual-Marqui, R. D., Milz, P., & Kochi, K. (2012). Reduced functional connectivity between cortical sources in five meditation traditions detected with lagged coherence using EEG tomography. Neuroimage, 60(2), 1574–1586. doi:10.1016/j.neuroimage.2012.01.042

Lomas, T., Ivtzan, I., & Fu, C. H. (2015). A systematic review of the neurophysiology of mindfulness on EEG oscillations. Neurosci Biobehav Rev, 57, 401–410. doi:10.1016/j.neubiorev.2015.09.018

Lueke, A., & Gibson, B. (2015). Mindfulness Meditation Reduces Implicit Age and Race Bias: The Role of Reduced Automaticity of Responding. Social Psychological and Personality Science, 6(3), 284–291. doi:10.1177/1948550614559651

Lutz, A., Dunne, J. D., & Davidson, R. J. (2006). Meditation and the Neuroscience of Consciousness: An Introduction. In The Cambridge Handbook of Consciousness.

Lutz, A., Greischar, L. L., Rawlings, N. B., Ricard, M., & Davidson, R. J. (2004). Long-term meditators self-induce high-amplitude gamma synchrony during mental practice. PNAS.

Mair, P., & Wilcox, R. (2020). Robust statistical methods in R using the WRS2 package. Behavior research methods, 52(2), 464. doi:10.3758/s13428-019-01246-w

Mathewson, K. E., Lleras, A., Beck, D. M., Fabiani, M., Ro, T., & Gratton, G. (2011). Pulsed out of awareness: EEG alpha oscillations represent a pulsed-inhibition of ongoing cortical processing. Frontiers in psychology, 2(MAY), 99. doi:10.3389/fpsyg.2011.00099

Oostenveld, R., Fries, P., & Maris, E. (2011). FieldTrip: Open Source Software for Advanced Analysis of MEG, EEG, and Invasive Electrophysiological Data. Computational Intelligence and Neuroscience : CIN, 2011(2011), 156869. doi:10.1155/2011/156869

Ouyang, G., Hildebrandt, A., Schmitz, F., & Herrmann, C. S. (2020). Decomposing alpha and 1/f brain activities reveals their differential associations with cognitive processing speed. Neuroimage, 205, 116304. doi:10.1016/j.neuroimage.2019.116304

Pang, D., & Ruch, W. (2019). Scrutinizing the Components of Mindfulness: Insights from Current, Past, and Non-meditators. Mindfulness, 10(3), 492–505. doi:10.1007/s12671-018-0990-4

Payne, J. R., Baell, O., Geddes, H., Fitzgibbon, B., Emonson, M., Hill, A. T., … Bailey, N. W. (2020). Experienced meditators exhibit no differences to demographically matched controls in theta phase synchronization, P200, or P300 during an auditory oddball task. Mindfulness, 11, 643–659.

Peterson, E., Rosen, B., Campbell, A., Belger, A., & Voytek, B. (2018). 1/f neural noise is a better predictor of schizophrenia than neural oscillations. BioRxiv. doi:10.1101/113449

Pion-Tonachini, L., Kreutz-Delgado, K., & Makeig, S. (2019). ICLabel: An automated electroencephalographic independent component classifier, dataset, and website. Neuroimage, 198, 181–197. doi:10.1016/j.neuroimage.2019.05.026

Pritchett, D. L., Siegle, J. H., Deister, C. A., & Moore, C. I. (2015). For things needing your attention: the role of neocortical gamma in sensory perception. Current Opinion in Neurobiology, 31, 254–263. doi:10.1016/j.conb.2015.02.004

Raichle, M. E., MacLeod, A. M., Snyder, A. Z., Powers, W. J., Gusnard, D. A., & Shulman, G. L. (2001). A default mode of brain function. Proc Natl Acad Sci U S A, 98(2), 676–682. doi:10.1073/pnas.98.2.676

Rihs, T., Michel, C., & Thut, G. (2009). A bias for posterior α-band power suppression versus enhancement during shifting versus maintenance of spatial attention. Neuroimage, 44(1), 190–199. doi:10.1016/j.neuroimage.2008.08.022

Robertson, M. M., Furlong, S., Voytek, B., Donoghue, T., Boettiger, C. A., & Sheridan, M. A. (2019). EEG power spectral slope differs by ADHD status and stimulant medication exposure in early childhood. Journal of neurophysiology, 122(6), 2427. doi:10.1152/jn.00388.2019

Rodriguez-Larios, J., Bracho Montes de Oca, E. A., & Alaerts, K. (2021). The EEG spectral properties of meditation and mind wandering differ between experienced meditators and novices. Neuroimage, 245, 118669. 10.1016/j.neuroimage.2021.118669

Rouder, J. N., Morey, R. D., Verhagen, J., Swagman, A. R., & Wagenmakers, E.-J. (2017). Bayesian Analysis of Factorial Designs. Psychological Methods, 22(2), 304–321. doi:10.1037/met0000057

Sauseng, P., Griesmayr, B., Freunberger, R., & Klimesch, W. (2010). Control mechanisms in working memory: A possible function of EEG theta oscillations. Neuroscience and Biobehavioral Reviews, 34(7), 1015–1022. doi:10.1016/j.neubiorev.2009.12.006

Somers, B., Francart, T., & Bertrand, A. (2018). Ageneric eeg artifact removal algorithm based on the multi-channel wiener filter. Journal of Neural Engineering, 15(3), 036007. doi:10.1088/1741-2552/aaac92

Steer, R. A., & Beck, A. T. (1997). Beck Anxiety Inventory.

Takahashi, T., Murata, T., Hamada, T., Omori, M., Kosaka, H., Kikuchi, M., … Wada, Y. (2005). Changes in EEG and autonomic nervous activity during meditation and their association with personality traits. Int J Psychophysiol, 55(2), 199–207. doi:10.1016/j.ijpsycho.2004.07.004

Tanaka, G. K., Peressutti, C., Teixeira, S., Cagy, M., Piedade, R., Nardi, A. E., … Velasques, B. (2014). Lower trait frontal theta activity in mindfulness meditators. Arq Neuropsiquiatr, 72(9), 687–693. doi:10.1590/0004-282X20140133

Taylor, V. A., Daneault, V., Grant, J., Scavone, G., Breton, E., Roffe-Vidal, S., … Beauregard, M. (2013). Impact of meditation training on the default mode network during a restful state. Soc Cogn Affect Neurosci, 8(1), 4–14. doi:10.1093/scan/nsr087

Voytek, B., & Knight, R. T. (2015). Dynamic Network Communication as a Unifying Neural Basis for Cognition, Development, Aging, and Disease. Biological Psychiatry, 77(12), 1089–1097. doi:10.1016/j.biopsych.2015.04.016

Voytek, B., Kramer, M. A., Case, J., Lepage, K. Q., Tempesta, Z. R., Knight, R. T., & Gazzaley, A. (2015). Age-Related Changes in 1/f Neural Electrophysiological Noise. J Neurosci, 35(38), 13257–13265. doi:10.1523/JNEUROSCI.2332-14.2015

Wang, M. Y., Freedman, G., Raj, K., Fitzgibbon, B. M., Sullivan, C. M., Tan, W. L., … Bailey, N. W. (2020). Mindfulness meditation alters neural activity underpinning working memory during tactile distraction. Cognitive, Affective, & Behavioral Neuroscience, 20(6), 1216–1233. doi:10.3758/s13415-020-00828-y

Whitten, T. A., Hughes, A. M., Dickson, C. T., & Caplan, J. B. (2011). A better oscillation detection method robustly extracts EEG rhythms across brain state changes: The human alpha rhythm as a test case. NeuroImage (Orlando, Fla.), 54(2), 860–874. doi:10.1016/j.neuroimage.2010.08.064

Wong, W. P., Camfield, D. A., Woods, W., Sarris, J., & Pipingas, A. (2015). Spectral power and functional connectivity changes during mindfulness meditation with eyes open: A magnetoencephalography (MEG) study in long-term meditators. International Journal of Psychophysiology, 98(1), 95–111. doi:10.1016/j.ijpsycho.2015.07.006

