## Supplementary Materials for "The mindful brain at rest: neural oscillations and aperiodic activity in experienced meditators"

#### eBOSC

eBOSC provides the ability to distinguish periods of EEG activity that show oscillations in specific frequencies from periods that do not show oscillations, enabling a comparison of not only oscillatory power, but also the time spent showing oscillations above the 1/f activity between groups (Haller et al., 2018; Kosciessa et al., 2020). In addition to measures of oscillatory power without 1/f activity and the 1/f components slope and intercept, the eBOSC algorithm allows analysis of peak oscillatory frequencies.

#### Epoch and channel data

Epoch and channel descriptive statistics were collected from the pre-processing stages and can be viewed in Table 1. The number of epochs included in the analysis exceeded the minimum number used by Kosciessa et al. (2020) for all participants (which was removal of no more than 15%), indicating that the eBOSC computations were valid for all participants.

The measure of bad channels for the eyes open condition violated the homogeneity of variance assumption, and so a robust Yuen's t test was conducted which revealed no differences between groups ($t(46.22^{a}) = 0.89^{a}$, $p = .379^{a}$, $BF_{01} = 0.59$). No differences were found between groups for any other variable within the epoch and channel data (all $p >$ 0.10). These data can be viewed in Fig. 1.

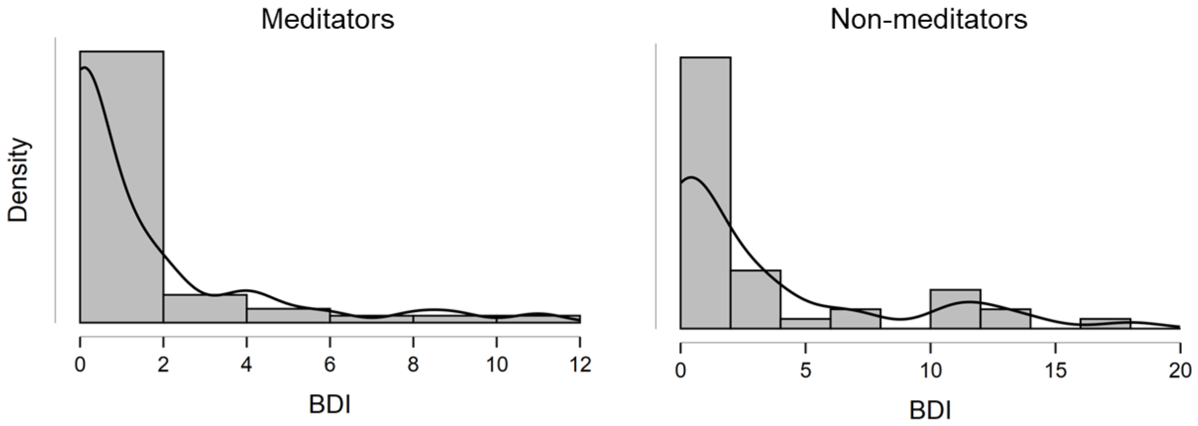


**Fig. 1** Violin plot demonstrating distribution and density overlay for BDI-II. These graphs demonstrate that majority of responses were zero or close to, with some degree of variation. Non-normal distribution was expected as demonstrated by the Bayes Factor value, as a score of 0 correlates with no symptoms associated with depression.

Table 1

*Descriptive Statistics for Epoch and Channel Data Acquired Through the Pre-Processing Stages*

| **Variable** | **Condition** | **Meditators** ***M (SD)*** *median and MAD used for robust tests* [a] | **Non-meditators** ***M (SD)*** *median and MAD used for robust tests* [a] | **$BF_{01}$** | **Frequentist statistics** |
|---|---|---|---|---|---|
| Bad epochs | Eyes open | 8.88 (7.17) | 9.23 (6.90) | 4.45 | $t(90) = 0.24$, $p = .811$ |
| Clean epochs | | 45.02 (8.80) | 42.64 (9.48) | 2.30 | $t(90) = -1.25$, $p = .214$ |
| Bad channels | | 2.5[a] (1.5[a]) | 3[a] (2[a]) | 1.70 | $t$(46.22[a]) = 0.89 [a], $p$ = .379[a] |
| Bad epochs | Eyes closed | 8.96 (6.7) | 7.11 (4.45) | 1.61 | $t(90) = -1.54$ $p = .127$ |
| Clean epochs | | 43.85 (8.88) | 43.77 (6.86) | 4.56 | $t(90) = -0.05$, $p = .961$ |
| Bad channels | | 2.92 (2.14) | 2.93 (2.04) | 4.57 | $t(90) = 0.04$, $p = .972$ |

*Note*. Frequentist statistics refer to student's t-test or robust Yuen's t tests. Yuen's t tests are indicated by a. The median and MAD are reported for robust tests. The mean and SD are reported for parametric tests. Robust tests are based on a trimmed sample and so the degrees of freedom (if given) will not be the same for other measures. BF = Bayes Factors. $BF_{01} > 1$ favours the model of the null hypothesis. $BF_{01}$ (BFexcl) is reported for non-significant findings whilst $BF_{10}$ (BFincl) is reported for significant findings. Higher-order interactions are excluded. All significance levels ($\alpha$) set at .05.

### RAGU

For statistical comparisons of each primary measure, a whole scalp analysis was conducted using Randomisation Graphical User Interface (RAGU) (Koenig et al. 2011). RAGU uses reference-free Root Mean Squared (RMS) measures and randomisation statistics to assess differences in the global neural response strength and topographical distributions across electrodes without a priori assumptions. RAGU also controls for multiple comparisons using permutation statistics (see Koenig, Kottlow, Stein, & Melie-García (2011)). RAGU allows for a combined comparison of the overall neural strength and distribution of activity using the Topographic Analysis of Variance (TANOVA), without normalising the amplitude of the data.

TANOVAs without L2 normalisation were run to test for an overall effect of differences between groups, as these analyses provide an indication of differences across both distribution and amplitude. RAGU also provides a t-map alongside the TANOVAs (for both without L2 normalisation and with L2 normalisation), with the t-min value produced indicating where meditators showed more negative values than non-meditators, and the t-max value indicating where meditators showed more positive values than non-meditators.

As the average reference was computed prior to the frequency transformed data, the typical global field power (GFP) test is converted to a comparison of the RMS, which is a valid indication of neural response strength in the frequency domain (Koenig et al., 2011).

### Frequentist analyses

Traditional frequentist statistical analyses were also conducted in R using the WRS2 package to implement robust statistics for mean peak frequency, RMS statistics for all oscillations measured, and the 1/f components (Mair & Wilcox, 2020; R Core Team, 2020). For any variables which demonstrated violations in homogeneity of variance, the robust Yuen's t-test was used instead of the independent samples t-test. See Tables 2, 3, 6 and 8 for further details and results of the tests conducted.

**Percentage of the EEG trace showing oscillatory activity**

The percentage of the EEG trace showing oscillatory activity reflects the oscillatory activity that is observed over the entire EEG recording and offers valuable insight as to which oscillatory frequencies are generated more commonly by participants across the time of the EEG recording. Whilst oscillatory power is an average of the whole EEG recording, the percentage of the EEG trace showing oscillatory activity can reflect the consistency of the oscillatory activity. The percentage of the EEG trace showing oscillatory activity is useful to examine in mindfulness meditators, as an average of oscillatory power could include extreme low or high power short bursts, which would be reflected as either low or high power, and not necessarily offer valuable insight as to the functional implications of oscillatory power.

Exploratory robust ANOVAs were conducted using the WRS2 statistics package in R to test for differences in the percentage of EEG trace showing oscillations in the frequency of interest for all oscillations (and for all electrodes).

There was a significant main effect found for the percentage of EEG trace showing theta oscillations, with meditators demonstrating a greater percentage of the EEG trace showing oscillations within the theta frequency compared to non-meditators (value = 5.11, $p = .026$). A full table of results from these analyses can be found in Table 6 and Table 7. No further differences were found between groups, and no interaction effects were present.

There was also a main effect of eyes open/closed for alpha and gamma oscillations. For alpha oscillations, the eyes closed condition demonstrated a greater percentage of the EEG trace showing oscillations within the alpha frequency compared to the eyes open condition (value = 28.26, $p < .001$), whilst for gamma oscillations, the eyes open condition demonstrated a greater percentage of the EEG trace showing gamma oscillations compared to the eyes closed condition (value = 6.1, $p = .016$). A full table of results from these analyses can be found in Table 6 and Table 7. No further effects were found (all $p > 0.05$).

### Mean peak frequency

The peak of alpha and theta frequencies may offer further insight as to the mechanisms of action behind mindfulness meditation (MM), specifically cognitive and attentional changes. Alpha peak frequency has been associated with information processing, increased task demands, sampling of sensory information, and the general inhibitory functions of alpha activity (Mierau, Klimesch, & Lefebvre, 2017). Theta peak frequency has been associated with similar functions of theta activity such as working memory, but there is limited research investigating its role, especially in meditation state- or trait-related contexts (Moran et al., 2010).

As mean peak frequency is a measure of the average number of oscillatory peaks per second, TANOVA or RMS analyses were not appropriate for these values. Exploratory robust ANOVAs were conducted using the WRS2 statistics package in R to test for differences in the mean peak frequency at electrodes Fz, PO7, and PO8 (electrodes that were selected as previous research suggested that theta and alpha power is maximal at these electrodes),

Differences in the mean peak frequency were found between groups for theta, alpha and beta mean peak frequency, with meditators demonstrating higher values for their peak frequencies in these oscillation ranges (except for alpha at Fz with eyes open, which showed a slower oscillatory frequency in the meditator group: Meditators Mdn (9.80); MAD (0.66), Non meditators Mdn (10.00); MAD (0.57)) albeit with small BF values (all $p < 0.05$, all BFincl > 0.857). Further information regarding these exploratory analyses can be found in Table 8.

### Exploratory findings of alpha peak frequency

The findings of the present study provide support of Saggar et al. (2015)'s model of alpha mean peak frequency, where the average number of oscillatory peaks per second was lower for meditators than non-meditators – demonstrating slower alpha cycles on average (although the present study was not a direct replication). Previous research has explored alpha peak frequency in a variety of task-related and state-related contexts (Mierau,

Klimesch, & Lefebvre, 2017). Although the role of alpha peak frequency is unclear, it is thought to be involved in information processing, increased task demands, and sampling of sensory information (Mierau et al., 2017). For the present study, meditators demonstrated a lower median peak alpha frequency compared to non-meditators for the electrodes Fz (in the eyes closed condition only), PO7 and PO8 (see Table 8). This same trend was also found by Saggar et al. (2015), where differences were most prominent at parieto-occipital electrodes, rather than frontal electrodes (and therefore frontal regions). Saggar et al. (2015) suggests that lower mean peak alpha frequency might arise from the effect of meditation on different cortical circuits and suggests a mechanistic neurophysiological explanation for the demonstrated effects of meditation on attention function.

Table 2

*RMS analysis for oscillatory power*

| **Variable** | **Condition** | **Meditators *Mdn* (*MAD*) (from R)** | **Non-meditators *Mdn* (*MAD*) (from R)** | **Meditators *Mean* (*SD*) (from R)** | **Non-meditators *Mean* (*SD*) (from R)** | **Effect/ interaction** | **Robust ANOVA (R)** | ***Confidence interval [CI] (from R)*** | ***Statistics from RAGU*** | ***Bayesian statistics*** |
|---|---|---|---|---|---|---|---|---|---|---|
| Alpha power (a.u.) | Eyes open | 483201.4 (298702.5) | 445314.1 (157038.5) | 572586.3 (327309.9) | 499929.1 (233485.5) | Main effect | value = 17.2, *p* < .001*** | [515626.4, 1481389] | ηp² = .057, *p* = .029* | BFincl = 1.462 |
| 41 m, 40 non m | Eyes closed | 2380160.7 (641271.7) | 1332889.2 (1062783.4) | 2170571 (997622.2) | 1592907 (1394519.3) | Condition | value = 121.37, *p* < .001*** | [2169790.7, 3135553] | ηp² = .591, *p* < .001*** | BFincl = 8.183e +16 |
| | | | | | | Interaction | value = 13.36, *p* < .001*** | [397090.3, 1362853] | ηp² = .048, *p* = .203 | BFincl = 1.473<br>BFexcl = 0.679 |
| Theta power (a.u.) | Eyes open | 333974.4 (344038.9) | 387094.6 (167288.2) | 686718.3 (649079) | 465947.6 (346273.6) | Main effect | value = 15.07, *p* < .001*** | [264201.57, 826111.0] | ηp² = .145, *p* < .001*** | BFincl = 52.911 |
| 39 med, 41 non med | Eyes closed | 755755.3 (357971.1) | 416944.7 (345125.5) | 842774.4 (448657.6) | 446144.1 (390934.2) | Condition | value = 2.5, *p* = .12 | [-59187.52, 502721.9] | ηp² = .014, *p* = .313 | BFexcl = 3.650 |
| | | | | | | Interaction | value = 2.93, *p* = .093 | [-40624.12, 521285.3] | ηp² = .025, *p* = .160 | BFexcl = 1.794 |
| Beta power (a.u.) | Eyes open | 67590.57 (24068.35) | 61969.95 (22093.08) | 70277.36 (24199.57) | 63505.48 (20724.44) | Main effect | value = 1.01, *p* = .319 | [-9461.65, 28732.50] | ηp² = .016, *p* = .288 | BFexcl = 1.991 |
| 37 med, 36 non med | Eyes closed | 97940.34 (31052.58) | 97101.56 (37683.77) | 103620.2 (28440.82) | 97499.09 (39376.26) | Condition | value = 49.87, *p* < .001*** | [48626.03, 86820.18] | ηp² = .636, *p* < .001*** | BFincl = 3.148e +14 |
| | | | | | | Interaction | value = 0.16, *p* = .689 | [-22953.98, 15240.17] | ηp² < .001, *p* = .947 | BFexcl = 4.212 |
| Gamma power (a.u.) | Eyes open | 7667.52 (3123.46) | 3817.31 (1294.39) | 8982.24 (4040.73) | 4358.78 (1965.76) | Main effect | value = 58.41, *p* < .001*** | [5420.07, 9224.88] | ηp² = .258, *p* < .001*** | BFincl = 77313.447 |
| 48 med, 44 non med | Eyes closed | 9701.58 (3664.48) | 5718.17 (2916.30) | 10117.35 (3691.63) | 6622.16 (4494.85) | Condition | value = 12.12, *p* < .001*** | [1433.40, 5238.20] | ηp² = .234, *p* < .001*** | BFincl = 4715.458 |
| | | | | | | Interaction | value = 0.23, *p* = .63 | [-2366.21, 1438.60] | ηp² = .031, *p* = .136 | BFexcl = 1.033 |

*Notes.* RMS refers to root-mean-square. a.u. = arbitrary units resulting from Morlet wavelet transform measures of power after the subtraction of the eBOSC modelled 1/f non-oscillatory activity. The mean and standard deviation (SD) are reported for parametric tests. Robust tests are based on a trimmed sample and so the degrees of freedom (if given) will not be the same for other measures. The 'value' for these variables reflects the test statistic. BF = Bayes Factors. $BF_{01}$ > 1 favours the model of the null hypothesis. $BF_{10}$ > 1 favours the model of the alternative hypothesis. $BF_{01}$ (BFexcl) is reported for non-significant findings whilst $BF_{10}$ (BFincl) is reported for significant findings. For the TANOVA with L2 normalisation, the BFincl value reflects a test of the interaction between Group and Electrode. Main effect compares meditators and non-meditators. Condition compares eyes closed vs eyes open conditions. Interaction compares the group (meditators and non-meditators) and condition (eyes closed and eyes open). ηp² = Partial eta squared. Adjusted p-values are specified by BH-p, and uncorrected p-values are also reported. Higher-order interactions are excluded. All significance levels ($\alpha$) set at .05. *p < .05. **p < .01. ***p < .001.

Table 3

*RMS analysis for 1/f components*

| **Variable** | **Condition** | **Meditators** ***Mdn (MAD)*** | **Non-meditators** ***Mdn (MAD)*** | **Meditators** ***Mean (SD) (from R)*** | **Non-meditators** ***Mean (SD)) (from R)*** | **Effect/ interaction** | ***Robust ANOVA (R)*** | ***Confidence intervals [CI] (from R)*** | ***Statistics from RAGU*** | ***Bayesian statistics*** |
|---|---|---|---|---|---|---|---|---|---|---|
| Slope | Eyes open | 1.27 (0.24) | 1.37 (0.21) | 1.30 (0.24) | 1.321(0.23) | Main effect | value = 0.71, *p* = .4 | [-0.22, 0.09] | $\eta p^2 = .002$, *p* = .684 | BFexcl = 2.453 |
| | Eyes closed | 1.47 (0.27) | 1.50 (0.24) | 1.48 (0.26) | 1.50 (0.23) | Condition | value = 17.83, *p* < .001*** | [0.17, 0.49] | $\eta p^2 = .585$, *p* < .001*** | BFincl = 6.326e +15 |
| | | | | | | Interaction | value = 0.15, *p* = .701 | [-0.12, 0.18] | $\eta p^2 < .001$, *p* = .929 | BFexcl = 4.226 |
| Intercept | Eyes open | 5.92 (0.39) | 5.91 (0.36) | 5.98 (0.40) | 5.93 (0.39) | Main effect | value = 0.89, *p* = .347 | [-0.13, 0.37] | $\eta p^2 = .007$, *p* = .415 | BFexcl = 1.816 |
| | Eyes closed | 6.40 (0.37) | 6.29 (0.45) | 6.39 (0.41) | 6.32 (0.44) | Condition | value = 38.16, *p* < .001*** | [0.52, 1.02] | $\eta p^2 = .736$, *p* < .001*** | BFincl = 3.142e +24 |
| | | | | | | Interaction | value = 0.1, *p* = .754 | [-0.21, 0.29] | $\eta p^2 = .003$, *p* = .804 | BFexcl = 4.292 |
| | | | | | | | value = 0.71, *p* = .4 | | | |

*Notes*. RMS refers to root-mean-square. The mean and standard deviation (SD) are reported for parametric tests. Robust tests are based on a trimmed sample and so the degrees of freedom (if given) will not be the same for other measures. The ‘value’ for these variables reflects the test statistic. BF = Bayes Factors. $BF_{01} > 1$ favours the model of the null hypothesis. $BF_{10} > 1$ favours the model of the alternative hypothesis. $BF_{01}$ (BFexcl) is reported for non-significant findings whilst $BF_{10}$ (BFincl) is reported for significant findings. Main effect compares meditators and non-meditators. Condition compares eyes closed vs eyes open conditions. Interaction compares the group (meditators and non-meditators) and condition (eyes closed and eyes open). $\eta p^2$ = Partial eta squared. Adjusted p-values are specified by BH-p, and uncorrected p-values are also reported. Higher-order interactions are excluded. All significance levels ($\alpha$) set at .05. *p < .05. **p < .01. ***p < .001.

Table 4

*TANOVA Results for the Comparisons 1/f slope and intercept*

| **Variable** | **Effect/ interaction** | **TANOVA without L2 normalisation (testing for overall effect including both amplitude and distribution effects)** | **TANOVA with L2 normalisation (testing for distribution effects independent of overall amplitude differences)** |
|---|---|---|---|
| Slope | Main effect | $\eta p^2 = .005$, $p = .61$, BH adjusted p = .66 | $\eta p^2 = .018$, $p = .09$ |
| | Condition | $\eta p^2 = .393$, $p < .001$***, BH adjusted p = .003** | $\eta p^2 = .013$, $p = .294$ |
| | Interaction | $\eta p^2 = .008$, $p = .715$, BH adjusted p = .715 | $\eta p^2 = .013$, $p = .270$ |
| Intercept | Main effect | $\eta p^2 = .009$, $p = .378$, BH adjusted p = .467 | $\eta p^2 = .020$, $p = .046$* |
| | Condition | $\eta p^2 = .643$, $p < .001$***, BH adjusted p = .003** | $\eta p^2 = .022$, $p = .025$* |
| | Interaction | $\eta p^2 = .010$, $p = .623$, BH adjusted p = .66 | $\eta p^2 = .023$, $p = .016$* |

*Notes.* Main effect compares meditators and non-meditators. Condition compares eyes closed vs eyes open conditions. Interaction compares the group (meditators and non-meditators) and condition (eyes closed and eyes open). $\eta p^2$ = Partial eta squared. Adjusted p-values are specified by BH-p, and uncorrected p-values are also reported. Higher-order interactions are excluded. All significance levels ($\alpha$) set at .05. *p < .05. **p < .01. ***p < .001.

Table 5

*TANOVA Results for the Comparisons of Power Without 1/f*

| **Variable** | **Effect/ interaction** | **TANOVA without L2 normalisation (testing for overall effect including both amplitude and distribution effects)** | **TANOVA with L2 normalisation (testing for distribution effects independent of overall amplitude differences)** |
|---|---|---|---|
| Alpha power (a.u.) | Main effect | $\eta p^2 = .066$, $p = .006$**, BH adjusted p = .014* | $\eta p^2 = .040$, $p < .001$*** |
| | Condition | $\eta p^2 = .532$, $p < .001$***, BH adjusted p = .003** | $\eta p^2 = .240$, $p < .001$*** |
| | Interaction | $\eta p^2 = .057$, $p = .122$, BH adjusted p = .2 | $\eta p^2 = .027$, $p = .131$ |
| Theta power (a.u.) | Main effect | $\eta p^2 = .139$, $p < .001$***, BH adjusted p = .003** | $\eta p^2 = .038$, $p < .001$*** |
| | Condition | $\eta p^2 = .029$, $p = .110$, BH adjusted p = .198 | $\eta p^2 = .045$, $p < .001$*** |
| | Interaction | $\eta p^2 = .036$, $p = .071$, BH adjusted p = .142 | $\eta p^2 = .050$, $p < .001$*** |
| Beta power (a.u.) | Main effect | $\eta p^2 = .030$, $p = .169$, BH adjusted p = .254 | $\eta p^2 = .026$, $p = .061$ |
| | Condition | $\eta p^2 = .499$, $p < .001$***, BH adjusted p = .003** | $\eta p^2 = .203$, $p < .001$*** |
| | Interaction | $\eta p^2 = .022$, $p = .389$, BH adjusted p = .467 | $\eta p^2 = .032$, $p = .062$ |
| Gamma power (a.u.) | Main effect | $\eta p^2 = .191$, $p < .001$***, BH adjusted p = .003** | $\eta p^2 = .087$, $p < .001$*** |
| | Condition | $\eta p^2 = .052$, $p = .004$**, BH adjusted p = .01* | $\eta p^2 = .042$, $p = .002$** |
| | Interaction | $\eta p^2 = .015$, $p = .231$, BH adjusted p = .32 | $\eta p^2 = .028$, $p = .033$* |

*Notes.* a.u. = arbitrary units resulting from Morlet wavelet transform measures of power after the subtraction of the eBOSC modelled 1/f non-oscillatory activity. Main effect compares meditators and non-meditators. Condition compares eyes closed vs eyes open conditions. Interaction compares the group (meditators and non-meditators) and condition (eyes closed and eyes open). $\eta p^2$ = Partial eta squared. Adjusted p-values are specified by BH-p, and uncorrected p-values are also reported. Higher-order interactions are excluded. All significance levels ($\alpha$) set at .05. *p < .05. **p < .01. ***p < .001.

Table 6

*RMS Results for the percentage of EEG trace showing oscillations in the frequency of interest*

| **Variable** | **Condition** | **Meditators *Mdn (MAD) 48*** | **Non-meditators *Mdn (MAD) 44*** | **Meditators *Mean (SD) (from R)*** | **Non-meditators *Mean (SD)) (from R)*** | **Effect/ interaction** | **Robust ANOVA (R)** | ***Confidence Intervals [CI] (from R)*** | ***Statistics (from RAGU)*** | ***Bayesian statistics*** |
|---|---|---|---|---|---|---|---|---|---|---|
| Alpha percentage | Eyes open | 71.34 (18.22) | 63.15 (14.22) | 64.38 (19.95) | 63.17 (15.30) | Main effect | value = 2.86, *p* = .095 | [-1.69, 21.04] | ηp² = .003, *p* = .605 | BFexcl = 3.042 |
| | Eyes closed | 84.19 (12.99) | 81.64 (18.86) | 78.16 (20.33) | 75.67 (19.17) | Condition | value = 28.26, *p* < .001*** | [19.07, 41.80] | ηp² = .398, *p* < .001*** | BFincl = 5.048e +8 |
| | | | | | | Interaction | value = 0.002, *p* = .959 | [-11.07, 11.66] | ηp² = .002, *p* = .078 | BFexcl = 4.147 |
| Theta percentage | Eyes open | 28.53 (11.32) | 24.81 (7.10) | 30.75 (12.64) | 26.32 (9.94) | Main effect | value = 5.11, *p* = .026* | [0.81, 12.46] | ηp² = .022, *p* = .156 | BFincl = 0.868 BFexcl = 1.151 |
| | Eyes closed | 28.10 (9.61) | 26.23 (6.63) | 31.30 (13.91) | 28.84 (10.41) | Condition | value = 0.5, *p* = .481 | [-3.75, 7.91] | ηp² = .058, *p* = .018* | BFincl = 1.856 BFexcl = 0.539 |
| | | | | | | Interaction | value = 0.66, *p* = .421 | [-8.20, 3.45] | ηp² = .026, *p* = .137 | BFexcl = 1.887 |
| Beta percentage | Eyes open | 47.13 (10.85) | 47.35 (12.48) | 47.78 (12.29) | 48.07 (10.82) | Main effect | value = 0, *p* = .997 | [-7.08, 7.11] | ηp² < .001, *p* = .971 | BFexcl = 2.500 |
| | Eyes closed | 49.66 (12.81) | 47.18 (12.15) | 49.10 (10.97) | 48.645 (9.76) | Condition | value = 0.69, *p* = .408 | [-4.11, 10.07] | ηp² = .021, *p* = .182 | BFexcl = 2.548 |
| | | | | | | Interaction | value = 0.13, *p* = .718 | [-5.80, 8.39] | ηp² = .003, *p* = .594 | BFexcl = 3.872 |
| Gamma percentage | Eyes open | 15.75 (5.33) | 16.52 (3.88) | 16 (5.38) | 16.52 (4.34) | Main effect | value = 1.04, *p* = .311 | [-4.62, 1.48] | ηp² = .004, *p* = .561 | BFexcl = 2.161 |
| | Eyes closed | 13.62 (4.62) | 14.32 (5.08) | 14.31 (5.25) | 14.947 (4.68) | Condition | value = 6.1, *p* = .016* | [-6.85, -0.75] | ηp² = .250, *p* < .001*** | BFincl = 34210.604 |
| | | | | | | Interaction | value = 0.03, *p* = .869 | [-3.31, 2.79] | ηp² < .001, *p* = .867 | BFexcl = 4.123 |

*Notes.* RMS refers to root-mean-square. The mean and standard deviation (SD) are reported for parametric tests. Robust tests are based on a trimmed sample and so the degrees of freedom (if given) will not be the same for other measures. The 'value' for these variables reflects the test statistic. BF = Bayes Factors. $BF_{01}$ > 1 favours the model of the null hypothesis. $BF_{10}$ > 1 favours the model of the alternative hypothesis. $BF_{01}$ (BFexcl) is reported for non-significant findings whilst $BF_{10}$ (BFincl) is reported for significant findings. Main effect compares meditators and non-meditators. Condition compares eyes closed vs eyes open conditions. Interaction compares the group (meditators and non-meditators) and condition (eyes closed and eyes open). ηp² = Partial eta squared. Adjusted p-values are specified by BH-p, and uncorrected p-values are also reported. Higher-order interactions are excluded. All significance levels ($\alpha$) set at .05. *p < .05. **p < .01. ***p < .001.

Table 7

*TANOVA Results for the percentage of EEG trace for the oscillation*

| **Variable** | **Effect/ interaction** | **Non normalised statistics** |
|---|---|---|
| Alpha percentage | Main effect | $\eta p^2 = .003$, $p = .658$ |
| | Condition | $\eta p^2 = .361$, $p < .001$*** |
| | Interaction | $\eta p^2 = .002$, $p = .873$ |
| Theta percentage | Main effect | $\eta p^2 = .021$, $p = .153$ |
| | Condition | $\eta p^2 = .050$, $p = .003$** |
| | Interaction | $\eta p^2 = .018$, $p = .153$ |
| Beta percentage | Main effect | $\eta p^2 = .005$, $p = .596$ |
| | Condition | $\eta p^2 = .022$, $p = .077$ |
| | Interaction | $\eta p^2 = .009$, $p = .480$ |
| Gamma percentage | Main effect | $\eta p^2 = .008$, $p = .486$ |
| | Condition | $\eta p^2 = .075$, $p < .001$*** |
| | Interaction | $\eta p^2 = .008$, $p = .767$ |

*Notes.* Main effect compares meditators and non-meditators. Condition compares eyes open and eyes closed conditions. Interaction compares the group (meditators and non-meditators) and condition (eyes open and eyes closed). Only non-normalised statistics are presented as they non-normalised statistics provide a test of the distribution and amplitude, as an analysis of both power and distribution separately is not a valuable way to analyse this measure, as we have no reason to expect that the percentage of EEG trace showing an oscillation would differ by region only after normalising for the percentage of the EEG activity showing the oscillation across all electrodes. Therefore, both the overall amount of the EEG activity showing the oscillation across all electrodes is informative, and whether particular areas show a longer amount of time with the oscillation, but whether the oscillation is present in a specific area for proportionally longer compared to all other electrodes in one group compared to the other is not very informative. Median and MAD are reported for all TANOVAs. $\eta p^2$ = Partial eta squared. a.u. = arbitrary unit. All significance levels ($\alpha$) set at .05. *p < .05. **p < .01. ***p < .001.

Table 8

*Robust Mixed ANOVA Results for mean peak frequency*

| Variable | Condition | Electrode | Meditators *Mdn* (*MAD*) 48 | Non-meditators *Mdn* (*MAD*) 44 | Effect/ interaction | Robust ANOVA (R) | *Bayesian statistics* |
|---|---|---|---|---|---|---|---|
| Alpha mean peak frequency | Eyes open | Fz | 10.15 (0.45) | 10.12 (0.55) | Electrode | value = 0.9, *p* = .64 | BFexcl = 4.103 |
| | Eyes closed | | 9.80 (0.66) | 10.00 (0.57) | Condition | value = 38.1, *p* < .001*** | BFincl = 7.596e +33 |
| | Eyes open | PO7 | 10.19 (0.41) | 10.26 (0.42) | Group | value = 11.52, *p* < .001*** | BFincl = 0.856 |
| | Eyes closed | | 9.75 (0.60) | 10.00 (0.61) | Electrode*Condition | value = 0.93, *p* = .63 | BFexcl = 7.208 |
| | Eyes open | PO8 | 10.21 (0.62) | 10.30 (0.45) | Electrode*Group | value = 0.006, *p* = .997 | BFexcl = 25.288 |
| | Eyes closed | | 9.77 (0.56) | 9.96 (0.51) | Condition*Group | value = 2.09, *p* = .15 | BFexcl = 1.380 |
| | | | | | Electrode*Condition*Group | value = 0.13, *p* = .936 | BFexcl = 9.152 |
| Theta mean peak frequency | Eyes open | Fz | 6.23 (0.28) | 6.23 (0.35) | Electrode | value = 2.04, *p* = .37 | BFexcl = 4.741 |
| | Eyes closed | | 6.15 (0.26) | 6.24 (0.36) | Condition | value = 0.67, *p* = .42 | BFexcl = 7.182 |
| | Eyes open | PO7 | 6.16 (0.22) | 6.30 (0.24) | Group | value = 10.87, *p* = .002** | BFincl = 1.008 |
| | Eyes closed | | 6.19 (0.33) | 6.25 (0.27) | Electrode*Condition | value = 0.3, *p* = .863 | BFexcl = 10.969 |
| | Eyes open | PO8 | 6.14 (0.30) | 6.22 (0.32) | Electrode*Group | value = 0.75, *p* = .687 | BFexcl = 11.928 |
| | Eyes closed | | 6.12 (0.25) | 6.20 (0.30) | Condition*Group | value = 0.7, *p* = .404 | BFexcl = 1.643 |
| | | | | | Electrode*Condition*Group | value = 1.65, *p* = .442 | BFexcl = 4.538 |
| Beta mean peak frequency | Eyes open | Fz | 19.82 (0.50) | 19.85 (0.73) | Electrode | value = 46.8, *p* < .001*** | BFincl = 1.498e +21 |
| | Eyes closed | | 19.72 (0.75) | 19.97 (0.71) | Condition | value = 0.062, *p* = .81 | BFexcl = 10.223 |
| | Eyes open | PO7 | 19.21 (0.56) | 19.51 (0.48) | Group | value = 11.96, *p* < .001*** | BFincl = 1.085 |
| | Eyes closed | | 19.22 (0.59) | 19.47 (0.76) | Electrode*Condition | value = 0.87, *p* = .65 | BFexcl = 3.873 |
| | Eyes open | PO8 | 19.24 (0.62) | 19.53 (0.54) | Electrode*Group | value = 1.53, *p* = .467 | BFexcl = 3.214 |
| | Eyes closed | | 19.27 (0.65) | 19.57 (0.78) | Condition*Group | value = 0.33, *p* = .566 | BFexcl = 5.839 |
| | | | | | Electrode*Condition*Group | value = 0.36, *p* = .837 | BFexcl =15.111 |
| Gamma mean peak frequency | Eyes open | Fz | 31.48 (1.71) | 32.27 (2.47) | Electrode | value = 80.8, *p* < .001*** | BFincl = 1.136e +19 |
| | Eyes closed | | 31.11 (1.81) | 32.04 (2.00) | Condition | value = 4.99, *p* = .027* | BFincl = 1.066 |
| | Eyes open | PO7 | 35.10 (4.90) | 34.96 (4.39) | Group | value = 0.29, *p* = .594 | BFexcl = 4.908 |
| | Eyes closed | | 33.56 (4.01) | 34.12 (4.44) | Electrode*Condition | value = 1.86, *p* = .399 | BFexcl = 17.372 |
| | Eyes open | PO8 | 34.31 (4.66) | 34.29 (3.71) | Electrode*Group | value = 1.17, *p* = .561 | BFexcl = 17.702 |
| | Eyes closed | | 32.95 (3.88) | 33.24 (3.06) | Condition*Group | value = 0.46, *p* = .499 | BFexcl = 1.674 |
| | | | | | Electrode*Condition*Group | value = 0.77, *p* = .682 | BFexcl = 8.003 |

*Notes*. The ‘value’ for these variables reflects the test statistic. Median and MAD are reported for robust ANOVAs. Robust statistics do not report any degrees of freedom since an adjusted critical value is used. BF = Bayes Factors. $BF_{01}$ > 1 favours the model of the null hypothesis. $BF_{10}$ > 1 favours the model of the alternative hypothesis. $BF_{01}$ (BFexcl) is reported for non-significant findings whilst $BF_{10}$ (BFincl) is reported for significant findings. Partial eta squared = $\eta p^2$. All significance levels ($\alpha$) set at .05. **p* < .05. ***p* < .01. ****p* < .001.

Table 9

*Post-hoc Factorial Bayesian ANOVA for alpha, theta and gamma power*

| | Group*Condition*Electrode factorial Bayesian ANOVA | Bayesian statistics (BFincl) |
|---|---|---|
| Alpha power | Group | 1.900 |
| | Condition*Group | 3.342e +41 |
| | Electrode*Group | 5.286e +20 |
| Theta power | Group | 34.308 |
| | Condition*Group | 2.722e +37 |
| | Electrode*Group | 5.324e +64 |
| Gamma power | Group | 12785.437 |
| | Condition*Group | 1.717 |
| | Electrode*Group | 2.169e +94 |

*Notes.* BF = Bayes Factors. $BF_{01}$ > 1 favours the model of the null hypothesis. $BF_{10}$ > 1 favours the model of the alternative hypothesis. $BF_{01}$ (BFexcl) is reported for non-significant findings whilst $BF_{10}$ (BFincl) is reported for significant findings.

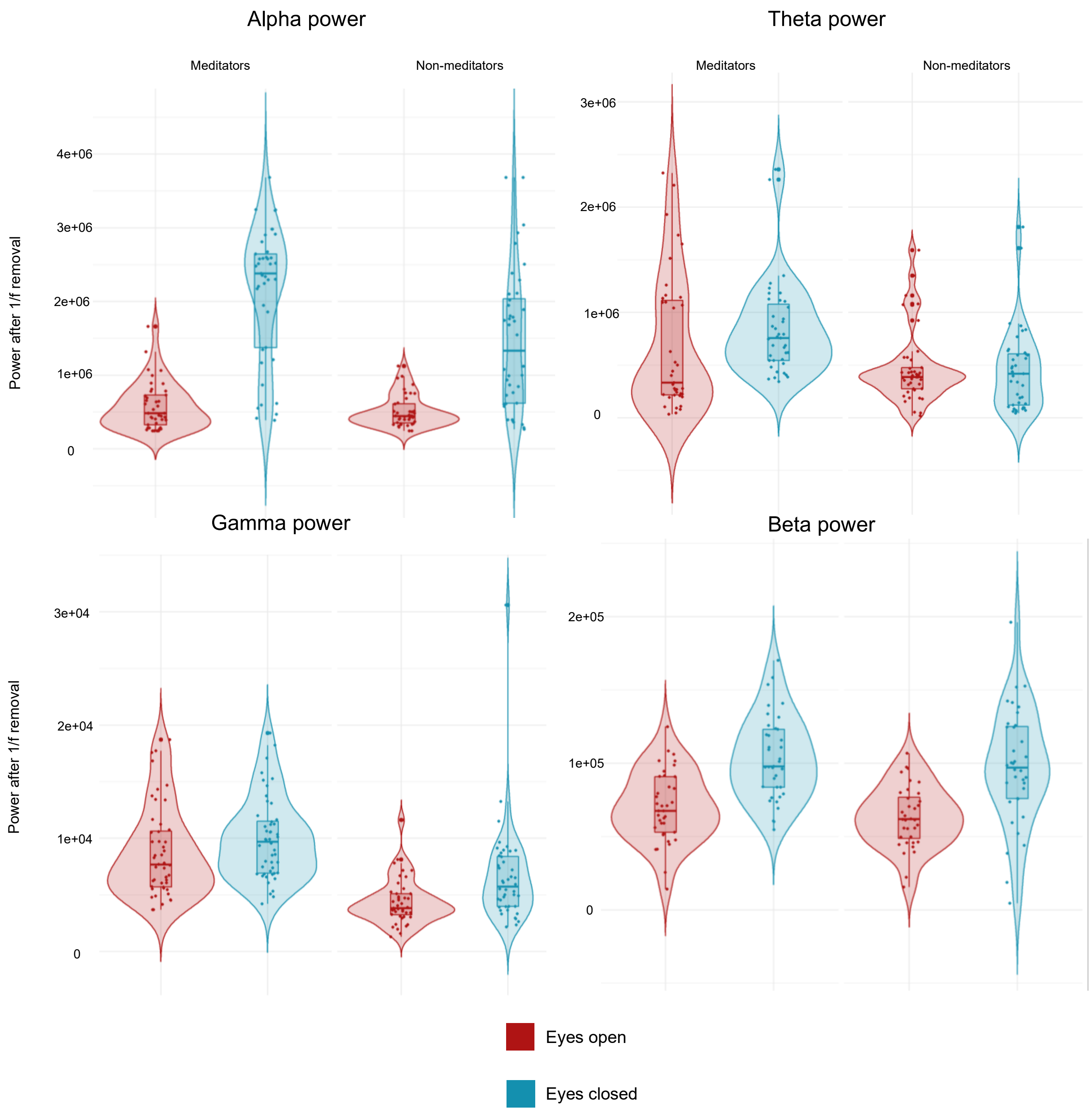


**Fig 2.** GFP violin plots for alpha, theta, gamma, and beta power after 1/f removal. This figure displays the GFP for all oscillations after 1/f removal. Interestingly, there is a greater spread – variance – for all eyes closed conditions for both meditators and non-meditators. Meditators appear to show a greater GFP values compared to non-meditators, except for alpha power.

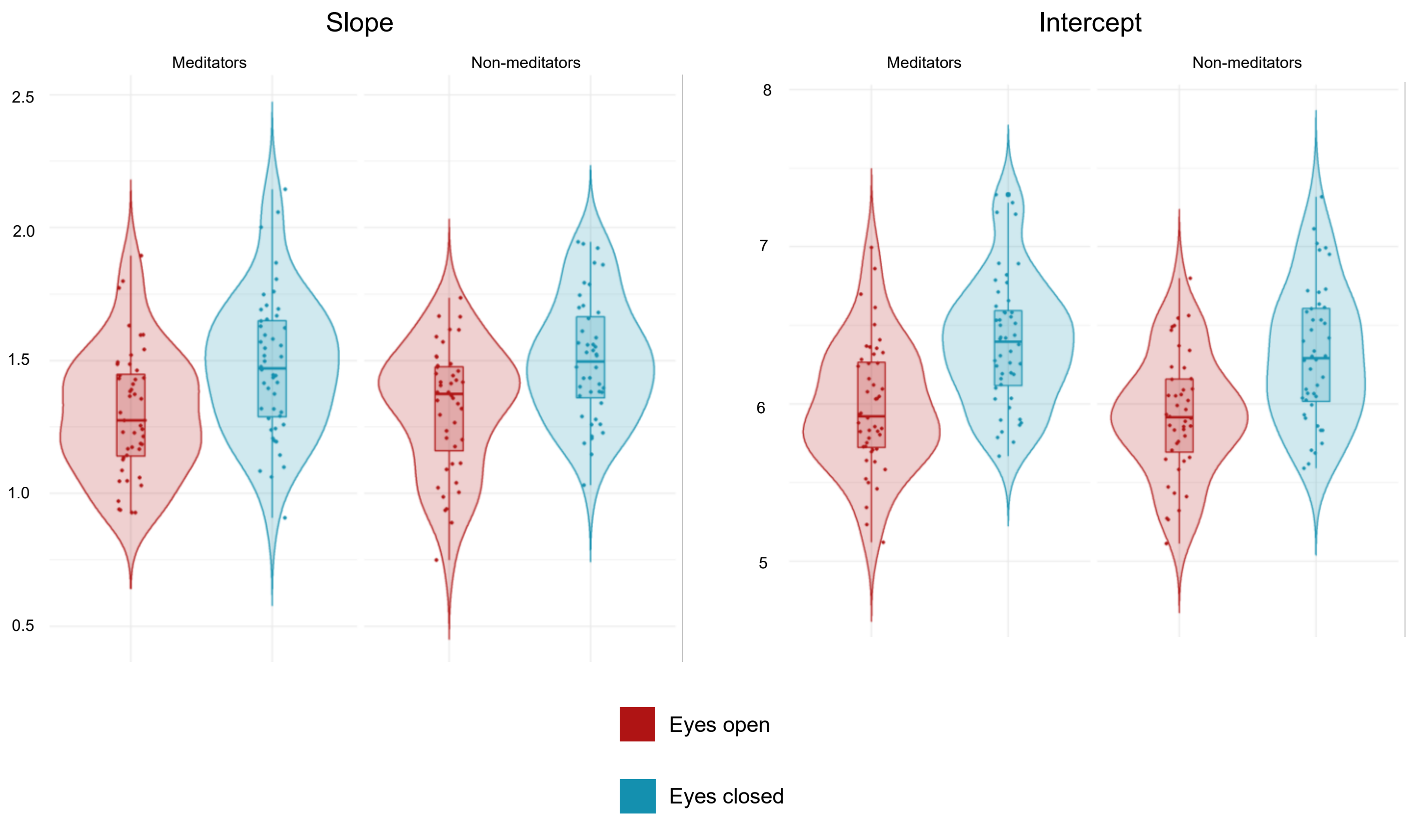


**Fig 3.** GFP violin plots for 1/f slope and intercept. This figure displays the GFP for 1/f slope and intercept. Both meditators and non-meditators, and for the eyes open and closed conditions demonstrate consistent values with a normal distribution and spread.

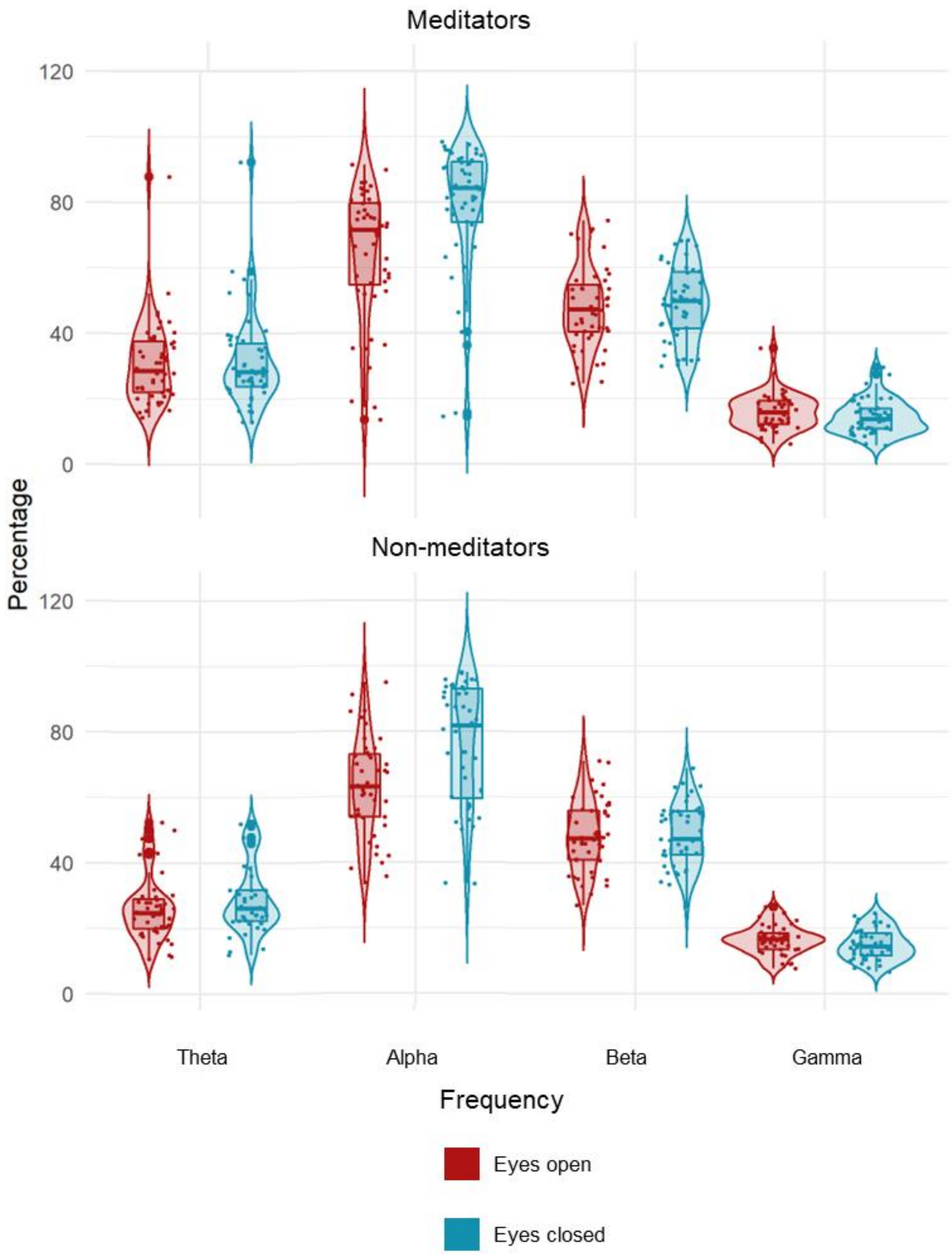


**Fig 4.** GFP violin plots for percentage of the EEG trace for alpha, beta, theta and gamma oscillations. This figure demonstrates how long each participant spent in each brain wave. Both meditators and non-meditators typically spent the most time in alpha waves, then beta, theta and a small percentage of time in gamma waves.

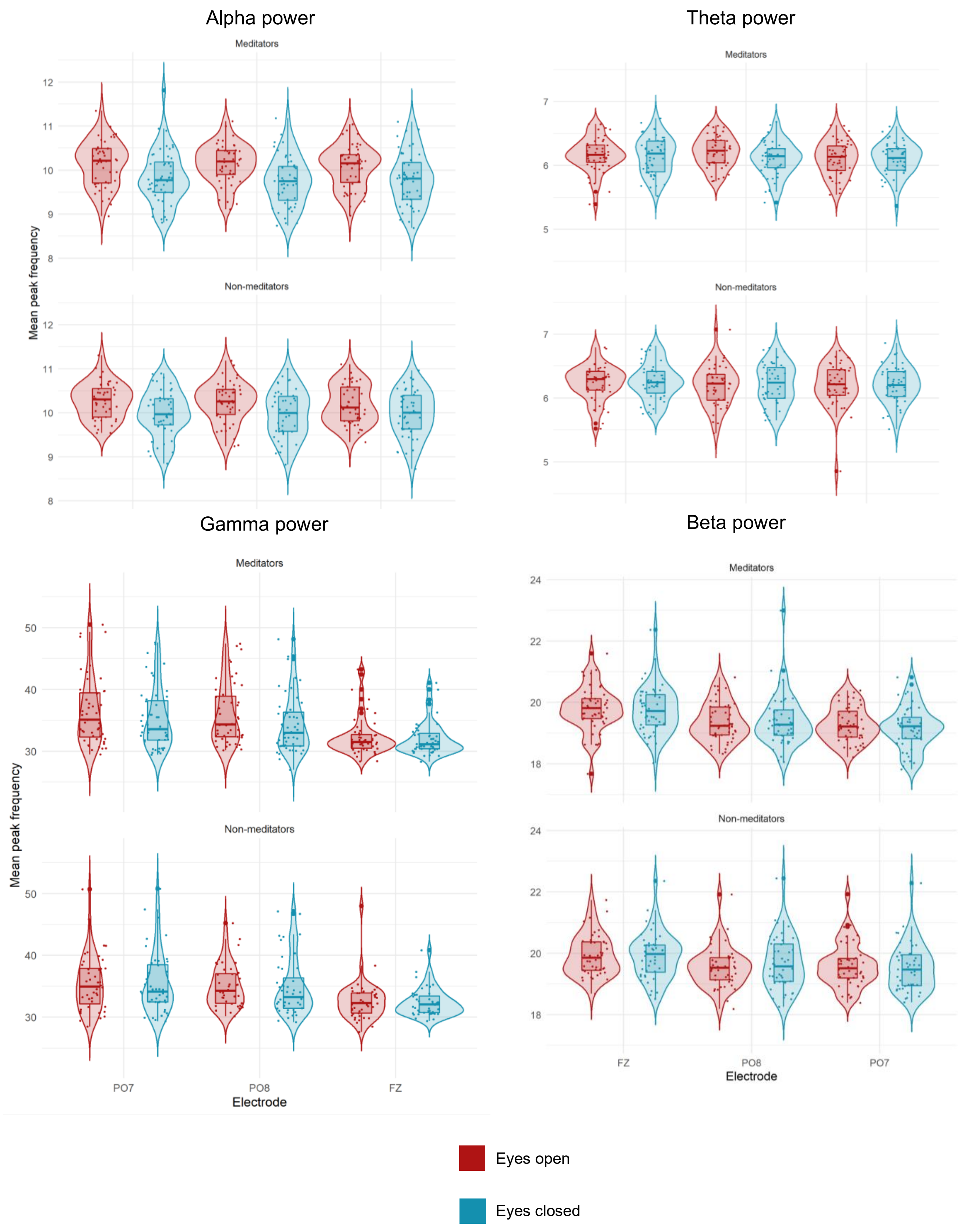


**Fig 5.** GFP violin plots for the mean peak of alpha, theta, gamma, and beta oscillations. These graphs demonstrate the mean peak of each oscillation. These graphs should be consistent as otherwise the results would not be a true reflection of each of these oscillations.

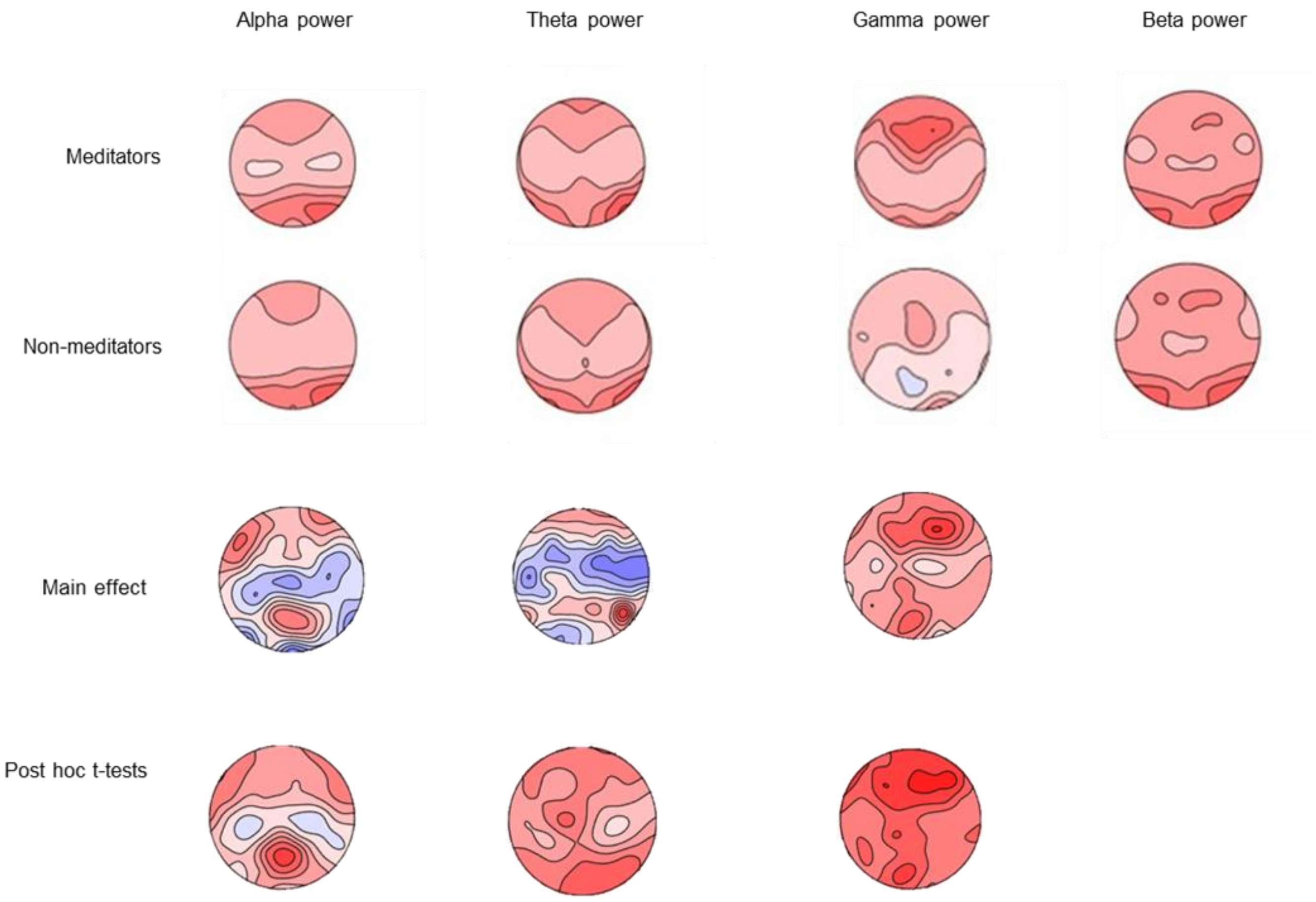


**Fig 6.** TANOVA graphs for each oscillatory power, significant interactions and main effects. This figure demonstrates the different GFP maps for each oscillation, significant main effects, and significant interactions.

### References

Haller, M., Donoghue, T., Peterson, E., Varma, P., Gao, R., Noto, T., . . . Voytek, B. (2018). Parameterizing neural power spectra. *BioRxiv*. doi:10.1101/299859

Koenig, T., Kottlow, M., Stein, M., & Melie-García, L. (2011). Ragu: A Free Tool for the Analysis of EEG and MEG Event-Related Scalp Field Data Using Global Randomization Statistics. *Computational Intelligence and Neuroscience, 2011*, 1-14. doi:10.1155/2011/938925

Kosciessa, J. Q., Grandy, T. H., Garrett, D. D., & Werkle-Bergner, M. (2020). Single-trial characterization of neural rhythms: Potential and challenges. *Neuroimage, 206*, 116331. doi:10.1016/j.neuroimage.2019.116331

Mair, P., & Wilcox, R. (2020). Robust statistical methods in R using the WRS2 package. *Behavior research methods, 52*(2), 464. doi:10.3758/s13428-019-01246-w

Moran, R., Campo, P., Maestú, F., Reilly, R., Dolan, R., & Strange, B. (2010). Peak Frequency in the Theta and Alpha Bands Correlates with Human Working Memory Capacity. Frontiers in human neuroscience, 4. doi:10.3389/fnhum.2010.00200

Mierau, A., Klimesch, W., & Lefebvre, J. (2017). State-dependent alpha peak frequency shifts: Experimental evidence, potential mechanisms and functional implications. *Neuroscience, 360*, 146-154. doi:10.1016/j.neuroscience.2017.07.037

Saggar, M., Zanesco, A. P., King, B. G., Bridwell, D. A., Maclean, K. A., Aichele, S. R., . . . Miikkulainen, R. (2015). Mean-field thalamocortical modeling of longitudinal EEG acquired during intensive meditation training. Neuroimage, 114, 88-104. doi:10.1016/j.neuroimage.2015.03.073
